# Spatial Clustering Analysis with Spectral Imaging-based Single-Step Multiplex Immunofluorescence (SISS-mIF) for Assisting Histological Diagnosis

**DOI:** 10.1101/2024.06.17.597874

**Authors:** Tomohiko Nakamura, Noe Kaneko, Towako Taguchi, Kenji Ikeda, Moe Sakata, Miori Inoue, Tetsuro Kuwayama, Hirokazu Tatsuta, Iichiroh Onishi, Morito Kurata, Kazuhiro Nakagawa

## Abstract

Precision medicine, based on spatial biology, is crucial for accurately diagnosing cancer and predicting drug responses. Here, we introduce the Spectral Imaging-based Single-Step Multiplex Immunofluorescence (SISS-mIF) technique, utilizing hyperspectral imaging to capture fluorescence spectra simultaneously. This approach optimizes tissue autofluorescence spectra for each image automatically, allowing the use of fluorescent direct-labeled antibodies for multicolor staining in a single step. Unlike conventional methods, the images are generated as standardized intensity independent of capture conditions, enabling consistent comparisons under different imaging conditions. This technique allows the detection of CD3, CD5, and CD7 in T-cell lymphoma on a single slide. The use of fluorescent direct-labeled antibodies enables triple staining of CD3, CD5, and CD7 without cross-reactivity, maintaining the same intensity as single stains. Furthermore, we developed a joint Non-Negative Matrix Factorization-based Spatial Clustering Analysis (jNMF-SCA) with a modified spectral unmixing system, highlighting its potential as a supportive diagnostic tool for T-cell lymphoma.

## Introduction

Immunohistochemistry (IHC) using 3,3’-diaminobenzidine (DAB)^1, 2^ serves as a foundational histochemical technique in routine pathology services for analyzing etiology and conducting histopathological diagnoses. However, detecting multiple biomarkers with conventional light microscopy is challenging, necessitating multiple serial tissue sections to ascertain the molecular and cellular locales of multiple biomarkers, significantly consuming tissue resources for examination and complicating the analysis of identical cells in small cells, such as lymphocytes. In diagnosing T-cell lymphoma, especially in cases with scattered infiltrative types of lymphoma cells, the depletion of T-cell surface markers such as CD3, CD5, or CD7 in tissue specimens indicates malignancy^3, 4^. However, confirming these biomarkers on the same cells for accurate diagnosis is difficult using conventional IHC with DAB. Although flow cytometry is the preferred diagnostic test technique, it is not universally accessible (Supplemental Table 1). The simultaneous detection of multiple antigens on histopathological sections has garnered interest for both research implications and for its utility in predicting the response to anticancer drugs by examining the spatial arrangement of molecules and cells within the tissue microenvironment^5^. Recently, various multiplex fluorescent immunostaining methods have been reported, each reviewed for its distinctive characteristics^6^.

The primary technique for identifying multiple biomarkers is fluorescence detection staining, which encompasses two principal categories. The first is a direct single-step staining method that employs only a primary antibody^7^ ^8^. The second approach involves multicolor staining and signal amplification through the sequential application of secondary antibodies, DNA oligos, enzymes, and other reactive molecules. The first category is represented by the direct method, while the second is represented by the indirect method^9^, the cycle staining method^10, 11^, the Tyramide Signal Amplification (TSA) method^12^, and the oligo barcode method^13–15^.

The direct method was the earliest approach to fluorescent immunostaining, relying on detecting the fluorescence intensity emitted by a dye attached to the antibody. However, its utility is limited by the challenge of distinguishing this fluorescence from the autofluorescence inherent in formalin-fixed paraffin-embedded (FFPE) tissue sections, a task complicated by the absence of signal amplification. As a result, the direct method has seen a decline in usage in contemporary practices. Conversely, the indirect method enhances the fluorescent signal by using a secondary antibody that binds to the primary antibody. However, this approach is limited by the specificity required in antibody combinations. The TSA method is capable of signal amplification through the covalent accumulation of fluorescently labeled tyramide molecules on a tissue section via an enzymatic reaction, but requires sequential antibody reactions. This process can obscure the target antigen with previously accumulated fluorescently labeled tyramide molecules, complicating subsequent antigen-antibody reactions and the analysis of co-expression within the same cell^6^. Cycle staining and oligo barcode methods aim to address these limitations by binding staining molecules followed by their desorption. However, the repetitive nature of staining, imaging, and desorption processes can cause tissue distortion and damage, potentially leading to misalignment in overlapping imaging data^6^.

Given these challenges, we aimed to refine the multiple fluorescent immunostaining method through the application of spectral separation technology developed for flow cytometry^16^ to tissue immunostaining. This methodological innovation is pivotal for obtaining spatial information through tissue immunostaining. Nonetheless, the emergence of wide-field imaging techniques, such as whole-slide imaging, complicates the identification of characteristic regions like the tissue microenvironment or scattered lymphoma cells. Existing methods that propose limiting the field of view to cells expressing specific biomarkers necessitate the establishment of thresholds, potentially introducing bias or overlooking significant regions^17^. To this effect, we developed a method that clusters samples based on the spatial distribution of cells in an unbiased manner across entire fields of view, enhancing our understanding of the tissue microenvironment and disease pathology beyond the capabilities of conventional methodologies.

## Results

### Spectral Imaging-based Single-Step Multiplex Immunofluorescence (SISS-mIF)

We employed direct single-step staining for a straightforward and rapid execution of multiple fluorescent immunostaining using multiple antibodies, each directly labeled with distinct fluorescent molecules, that react in a singular step with an antigen on a tissue specimen (Fig. 1a). Fluorescence spectral data are gathered at a consistent speed for each excitation via a line scan, mirroring the process of a bright-field digital pathology slide scanner (Fig. 1b, Supplemental Fig. 1). The fluorescence spectrum data is compiled into a matrix, with the spectrum organized as rows and the spatial dimensions of x, y as columns. In direct single-step staining, differentiating labeled fluorescence from autofluorescence is critical, given the finite number of fluorescent molecules that can be conjugated with antibodies, and autofluorescence spectra derived as components from serial sections. Unmixed images are procured by segregating the spectral data matrix, collected from the immunostained section, using the least squares method (LSM) with both optimized autofluorescence spectra from the spectral data matrix derived from the non-immunostained serial section through non-negative matrix factorization (NMF) ^18–23^ and the spectra of the fluorescence tagged to the antibodies (Fig. 1c).

**Figure 1.**
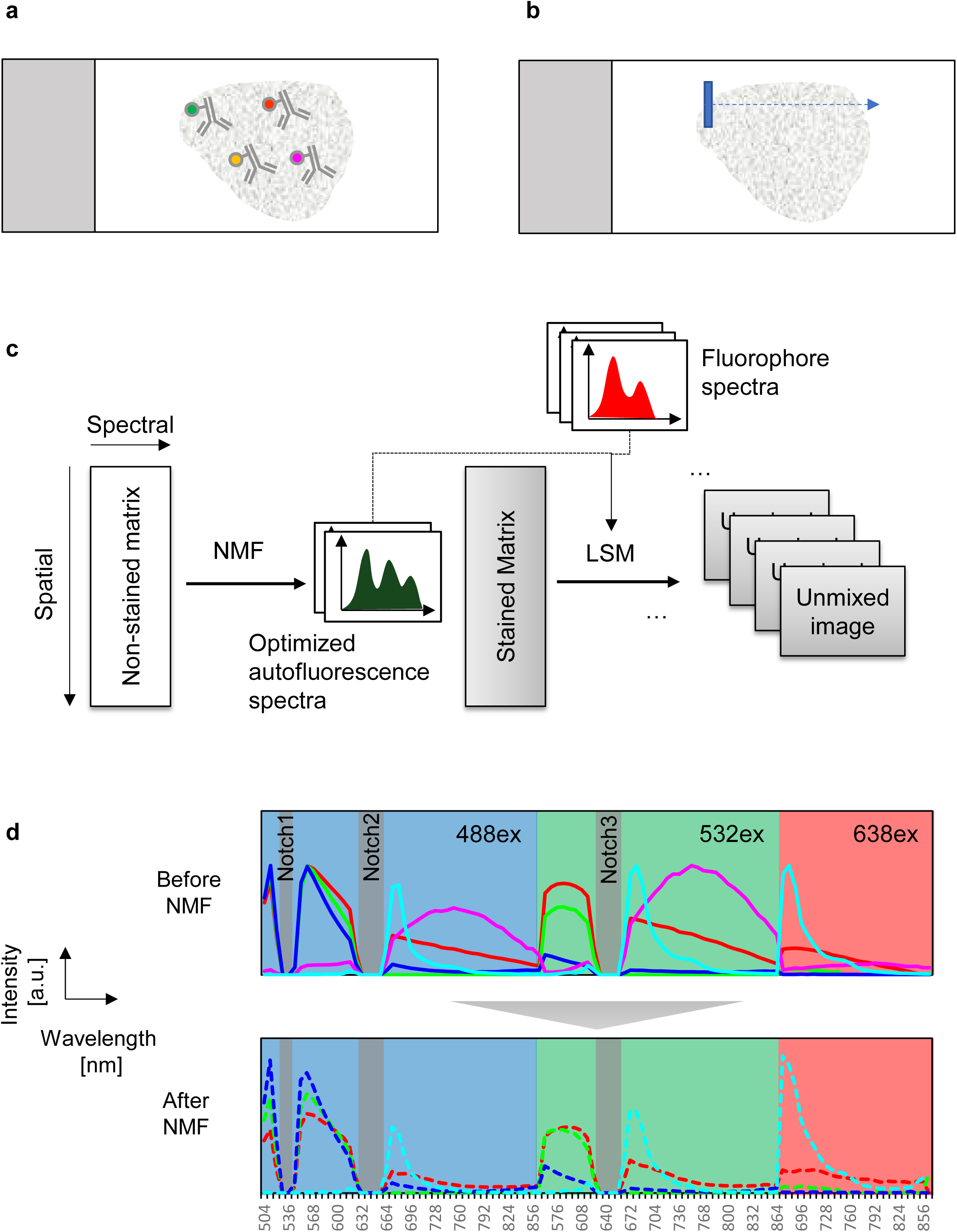
Summary of Spectral Imaging-based Single-Step Staining Multiplex Immunofluorescence (SISS-mIF). (a) Schematic depiction of multiplex direct single-step staining where tissue slides are stained with all fluorophore-conjugated primary antibodies simultaneously. (b) Illustration of spatial sweep spectroscopy, where three-dimensional spectral data are acquired by scanning perpendicular to the line, capturing the fluorescent spectrum of pixels. (c) Overview of image processing, highlighting autofluorescence component spectra extraction via non-negative matrix factorization (NMF) using an unstained slide, and subsequent least square method (LSM) unmixing with the stained matrix from the stained image, incorporating the standard spectrum including extracted autofluorescence spectra. (d) Spectra before and after optimization via NMF (520–552 nm, 624–664 nm are optical notch). Solid lines in the top chart represent five initial autofluorescence spectra inputted into the NMF algorithm (Red: Arachidonic Acid, Lime: Elastin, Blue: FAD, Magenta: PLD, Cyan: Protoporphyrin), while dashed lines in the bottom chart show optimized spectra by the NMF algorithm (Red: Auto Fluorescence1, Lime: Auto Fluorescence2, Blue: Auto Fluorescence3, Cyan: Auto Fluorescence4). The horizontal axis denotes wavelength [nm], and the vertical axis represents intensity [a.u.]. Fluorescence spectra excited by 4-lasers are concatenated along the wavelength direction.

The spectral unmixing process benefits from the acquisition of autofluorescence spectra, characterized by the separation into autofluorescence components and optimization of each component’s spectrum from actual sample data. The autofluorescence spectra from different cells vary based on the contained autofluorescence molecules^24^ and are also influenced by tissue fixation conditions^25, 26^. The separation of autofluorescence spectrum components aligns with cellular type differences and is optimized from spectral data acquired from unstained slides, starting with five fluorescence spectra: NADH and FAD as fluorescent nucleotides, arachidonic acid for lipids, porphyrins constituting hemoglobin and cytochrome, and the mounting agent (Fig. 1d). Each component spectrum’s optimization is executed via NMF, ensuring each spectrum is refined under conditions that preclude negative values to minimize residuals across all pixel spectral data^27, 28^ (Supplemental Fig. 2a). Unmixing of the original spectrum cube is performed by LSM using both the optimized autofluorescence spectra and the fluorescent antibody spectra^16^. The reference spectra for fluorescent antibodies are ascertained by slide samples wherein they are dispersed to a known concentration and normalized from the sensor surface volume as the concentration with one molecule present per unit voxel. The normalized reference spectra are fed into the LSM, generating standardized intensity independent of capture conditions as the molecules of equivalent soluble fluorophore (Supplemental Fig. 2b).

### Validation of SISS-mIF Image Processing

To validate image processing from autofluorescence spectrum extraction by NMF to that using spectrum separation by LSM, an evaluation was conducted using the image superimposed by simulation image and autofluorescence image (Fig. 2a). As a simulation image, spectral cube data with variance in intensity was generated considering shot noise and image sensor noise. As an autofluorescence image, an unstained sample was acquired using a hyperspectral imaging apparatus and superimposed on the simulation image. In superimposed images, the signal of the simulation image is buried in the autofluorescence signal, but this image processing separates the signal of the simulation image from the autofluorescence signal (Fig. 2b). In addition, image processing with unstained tissue images without superimposed simulation images correctly separated autofluorescence signals into the autofluorescence channel (Fig. 2c).

**Figure 2.**
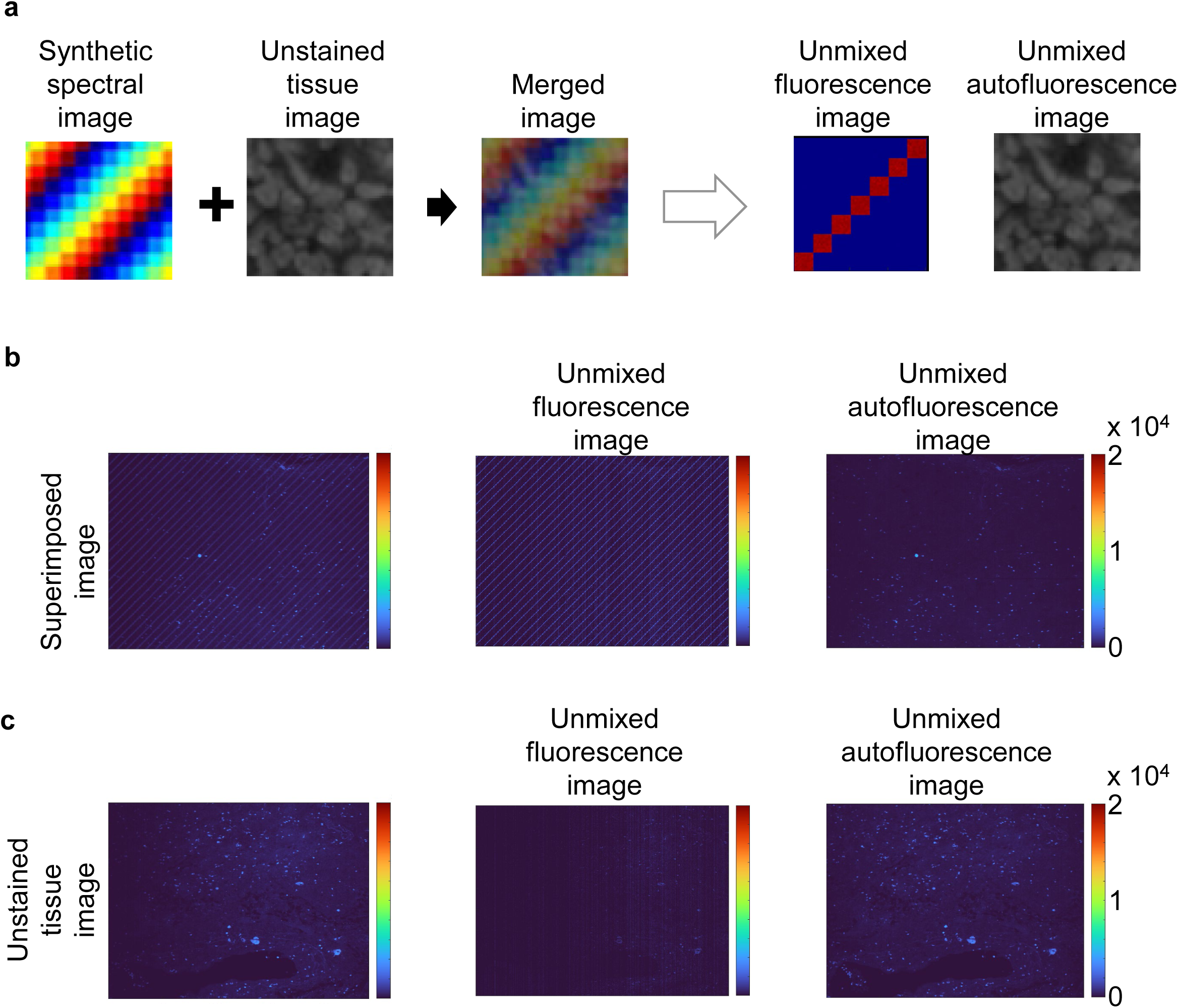
Validation of SISS-mIF Image Processing. (a) Schematic outline of generation of superimposed simulation image with unstained tissue image and spectral unmixing with SISS image processing. (b) Superimposed simulation image with unstained tissue image (left), fluorescence channel image (middle), and autofluorescence channel image (right) unmixed from superimposed simulation image with unstained tissue image. (c) Unstained tissue image (left), fluorescence channel image (middle), and autofluorescence channel image (right) unmixed from unstained tissue image.

### Direct Single-Step Staining Multiplex Immunofluorescence

Direct single-step staining for multiplex immunofluorescence was performed with three panels of fluorescently labeled antibodies. The first panel, designed for tonsil specimens, included Alexa Fluor (AF)488-CD4, AF555-PD-1, AF568-ki67, AF647-PDL1, AF680-CD3, AF700-Cytokeratin, AF750-CD8, and AF790-CD68, the second panel for skin comprised AF488-CD45RO, AF555-CD3, AF647-PD-1, AF680-Sox10, and AF750-Cytokeratin, and the third panel for breast contained AF488-CD4, eF570-Foxp3, eF615-CD20, AF647-CD8, AF680-CD68, and AF750-Cytokeratin, all showing specific staining without background for the majority of the markers (Fig. 3a–c). Since CD3, CD4, CD8, CD20, CD45RO, PD-1, and PD-L1 exhibited cell surface staining, ki67, Foxp3, and Sox10 demonstrated nuclear staining, and Cytokeratin and CD68 showed cytoplasmic staining, the determination of specific staining was based on the observed staining morphology and localization. As judged by three board certified pathologists, SISS-mIF showed similar staining patterns to the gold standard IHC (Supplemental Fig. 3).

**Figure 3.**
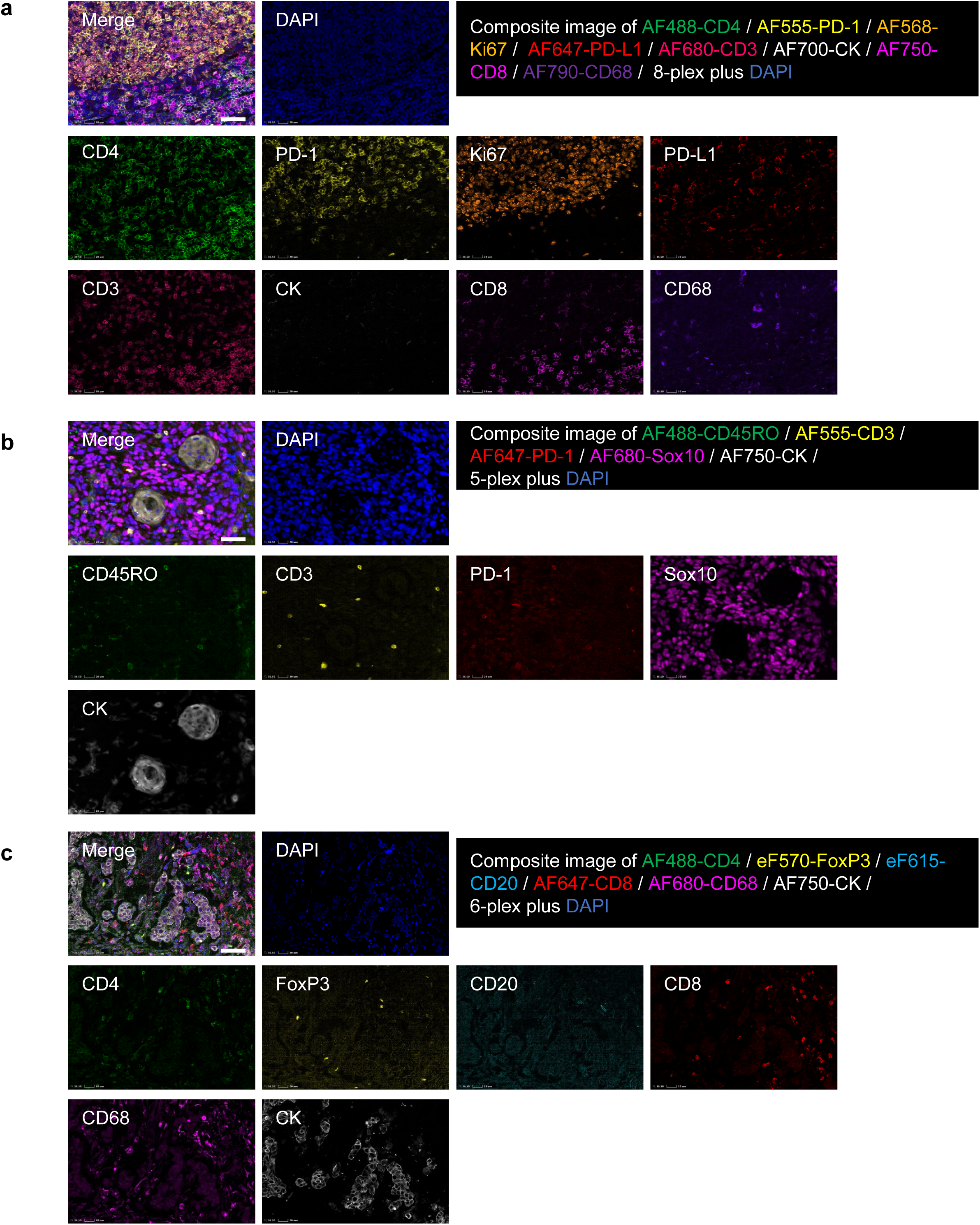
Direct Single-Step Staining Multiplex Immunofluorescence. (a) Representative 8-plex plus DAPI staining images on human tonsil tissue with merged and composite images showing AF488-CD4 in green, AF555-PD-1 in yellow, AF568-Ki67 in orange, AF647-PD-L1 in red, AF680-CD3 in pink, AF700-CK in white, AF750-CD8 in red-purple, AF790-CD68 in violet, and DAPI in blue. (b) Representative 5-plex plus DAPI staining images on human skin tissue with merged and composite images of AF488-CD45RO in green, AF555-CD3 in yellow, AF647-PD-1 in red, AF680-Sox10 in red-purple, AF750-CK in white, and DAPI in blue. (c) Representative 6-plex plus DAPI staining images on human breast tissue with merged and composite images of AF488-CD4 in green, eFluor (eF) 570-FoxP3 in yellow, eF615-CD20 in light blue, AF647-CD8 in red, AF680-CD68 in red-purple, AF750-CK in white, and DAPI in blue. Scale bars in all images are 50 µm. Deconvolution was applied to all images for image processing.

### Image Output in Standardized Intensity

In the evaluation of standardized intensity independent of capture conditions, images from slides stained under identical conditions from serial sections of a human tonsil tissue FFPE block were captured under varying scan conditions (Fig. 4a), and the resultant standardized intensities were juxtaposed (Fig. 4b). Comparable values were observed for all antibodies (AF488-αCD7, AF555-αCD3, and AF647-αCD5) under three different scan conditions: the first with standard settings, the second where the exposure time was halved, and the third where both the exposure time and the gain were reduced by half. In exploring the potential for co-expression analysis on the same cell, the standardized intensities following single, duplex, and triplex staining with α CD3, α CD3 and α CD5, and α CD3, α CD5, and α CD7 respectively on serial sections of human tonsil tissue were compared (Fig. 4c). The percentage of α-CD3 antibody in both duplex and triplex staining configurations was 97 ± 16% and 92 ± 10%, respectively, assuming the count in simplex staining as 100%. These findings indicate equivalent antibody presence across all staining modalities.

**Figure 4.**
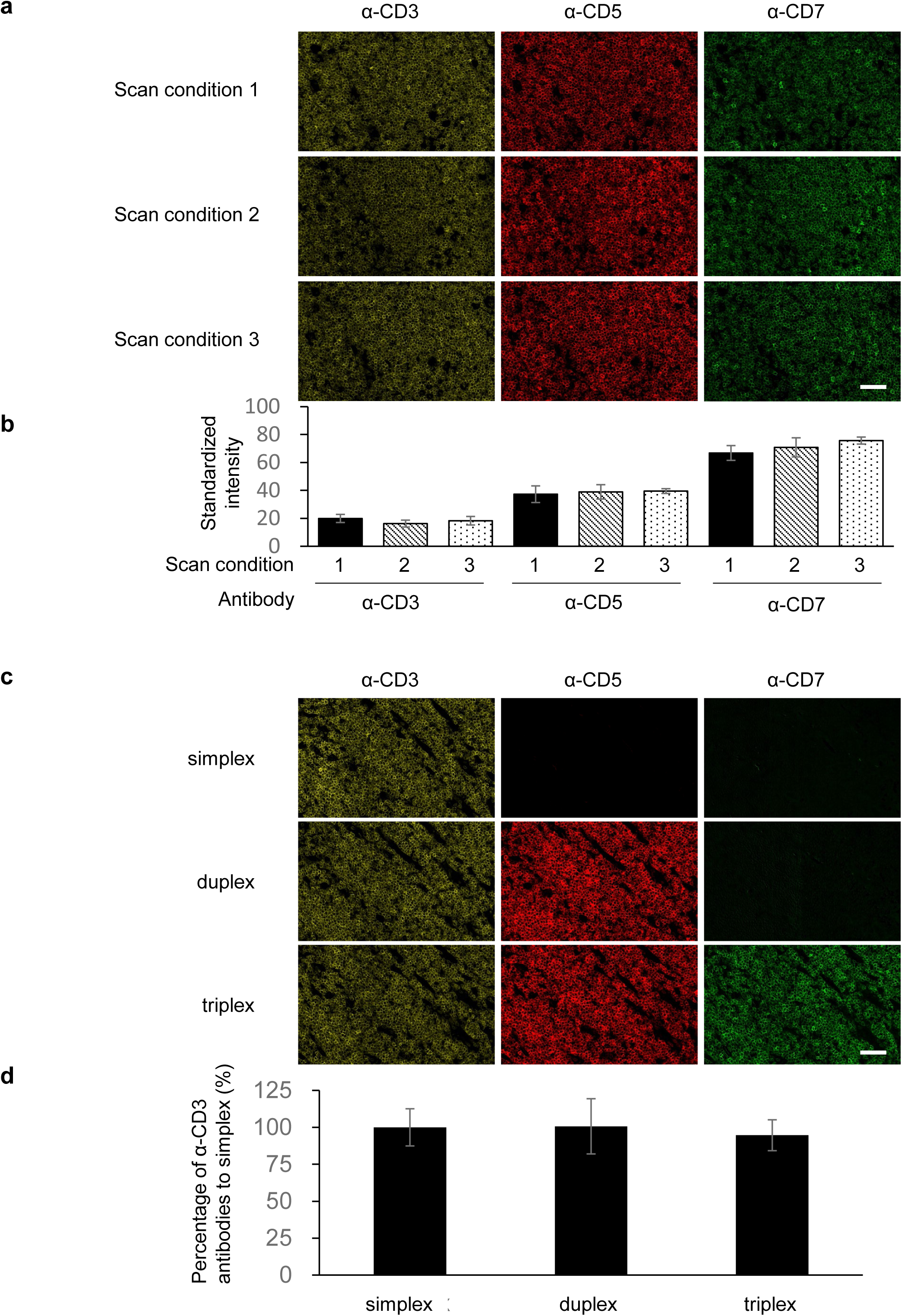
Image Outputs in Standardized Intensity. (a) Representative images captured under various exposure conditions and gain settings, with exposure condition and gain setting in scan condition 1 at 1800 µs and 9 dB, respectively. In scan condition 2, exposure conditions are set to 900 µs, and in scan condition 3, both exposure conditions and the gain are set to 900 µs and 3 dB. Multiplex staining for CD7 (green), CD3 (yellow), and CD5 (red) was performed on serial sections of human tonsils. Scale bars in all images are 50 µm. (b) Graph depicting standardized intensities under different exposure conditions, with error bars representing mean ± 1 SD for n = 3. (c) Staining of simplex, duplex, and triplex configurations performed with α-CD3, α-CD3 and α-CD5, α-CD3, α-CD5, and α-CD7, respectively, on serial sections of human tonsil. Scale bars in all images are 50 µm. (d) Graph showing α-CD3 standardized intensities in simplex, duplex, and triplex staining configurations, with error bars indicating mean ± 1 SD for n = 3. Deconvolution was applied to all images for image processing.

### Detection of CD3, CD5, and CD7 from Single Sections in T-cell Lymphoma

The efficacy of detecting CD3, CD5, and CD7 from a single section in T-cell lymphoma was evaluated by comparing the expression levels of each marker using DAB-IHC (IHC) and multiplex immunofluorescence (mIF) against their counterparts measured by flow cytometry (FCM). Instances of weak expression (e.g., FCM dim+, IHC weakly+) or absence detected by FCM, DAB-IHC, or mIF were deemed negative. Representative images of IHC and mIF staining are shown in Fig. 5 a–c and a comparison of FCM, IHC, and mIF results was tabulated (supplemental Table 2). An evaluation of CD3, CD5, and CD7 expression showed a concordance rate with FCM in 21 cases (84.0%) and 4 cases (16.0%) of discordance when utilizing mIF. Conversely, IHC indicated 18 cases (72.0%) of concordance and 7 cases (28.0%) of discordance. Among the six cases with CD5 deletions, mIF demonstrated a concordance in 5 cases (83.3%) and discordance in 1 case (16.7%), whereas IHC presented a 50-50 split with 3 concordant cases and 3 discordant cases. Similarly, in evaluating 18 patients with CD7 deletions, mIF achieved a concordance in 16 cases (88.9%) and discordance in 2 cases (11.1%), compared to IHC which showed 14 cases (77.8%) of concordance and 4 cases (22.2%) of discordance. Hence, mIF outperformed IHC across all metrics. For cases with discrepancies, mIF tended to be marginally more sensitive, with IHC-negative cases more likely to test positive with mIF (Table 1).

**Figure 5.**
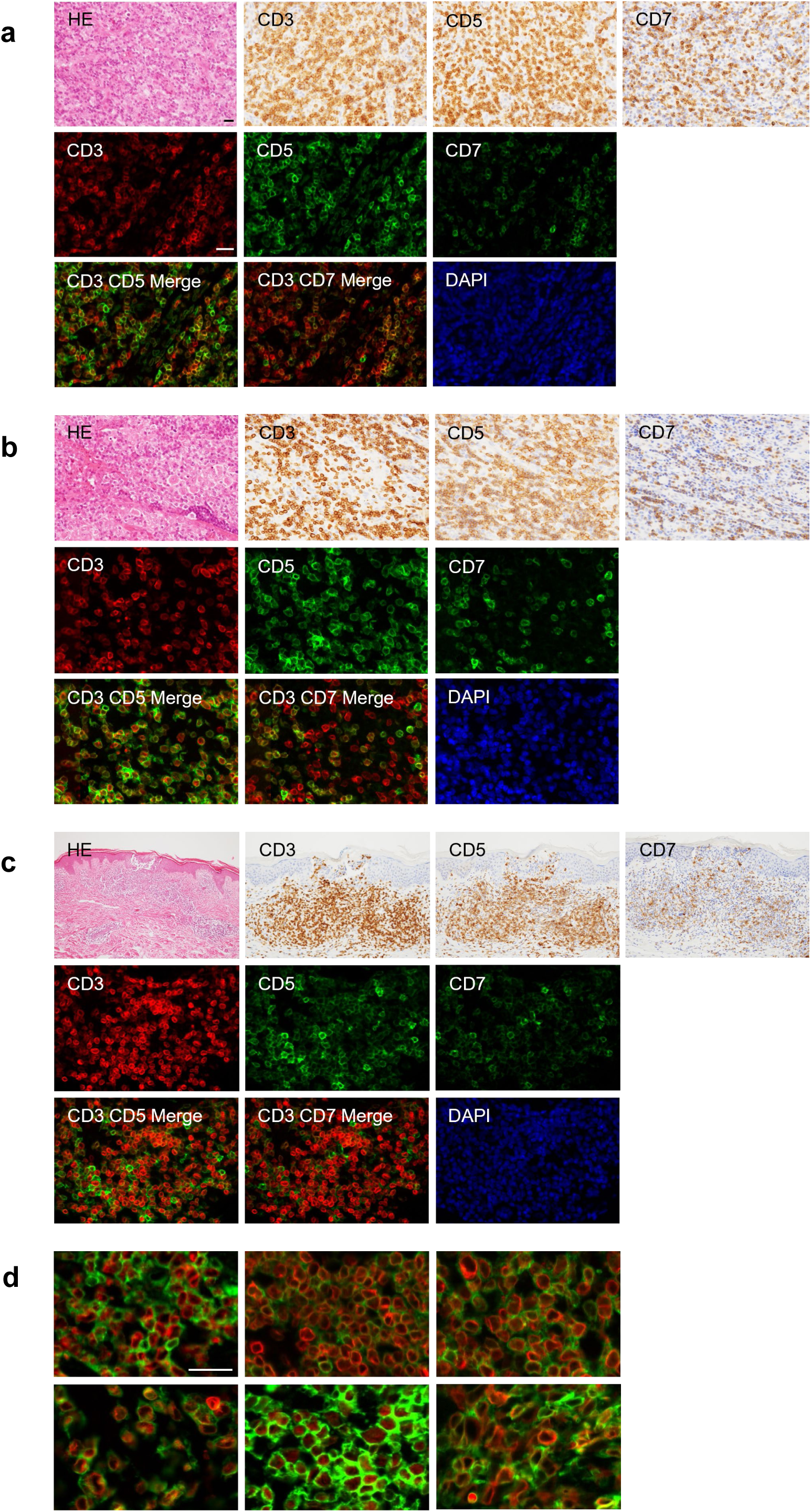
Detection of CD3, CD5, and CD7 in T-cell Lymphoma. (a–c) Representative histological, immunohistochemical (IHC), and multiplex immunofluorescence (mIF) images. (a) A lymph node diagnosed with angioimmunoblastic T-cell lymphoma (AITL), showing IHC results of CD3(+), CD5(+), and CD7(-) in a mixed population of normal and small T-cells, and corresponding mIF results. Flow cytometry (FCM) results (not displayed) are CD3(-), CD5(+), and CD7(-). (b) A lymph node with an AITL diagnosis, illustrating similar findings in both IHC and mIF results as CD3(+), CD5(+), and CD7(-), with FCM results (not shown) mirroring the pattern of CD3(-), CD5(+), and CD7(-). (c) A skin sample diagnosed with peripheral T-cell lymphoma (PTCL), showing IHC results of CD3(+), CD5(+), and CD7(-) ranging from negative to weakly positive, and mIF results showing CD3(+), CD5(+), and CD7(-) with dim positive expression. FCM results (not shown) show CD3(-), CD5(+), and CD7(-) with dim positive expression. (d) Representative images of CD3 cytoplasmic expression observed through mIF, offering a detailed visualization of intracellular protein localization. Scale bars in all images are set at 20 µm to facilitate comparison.

**Table 1.**
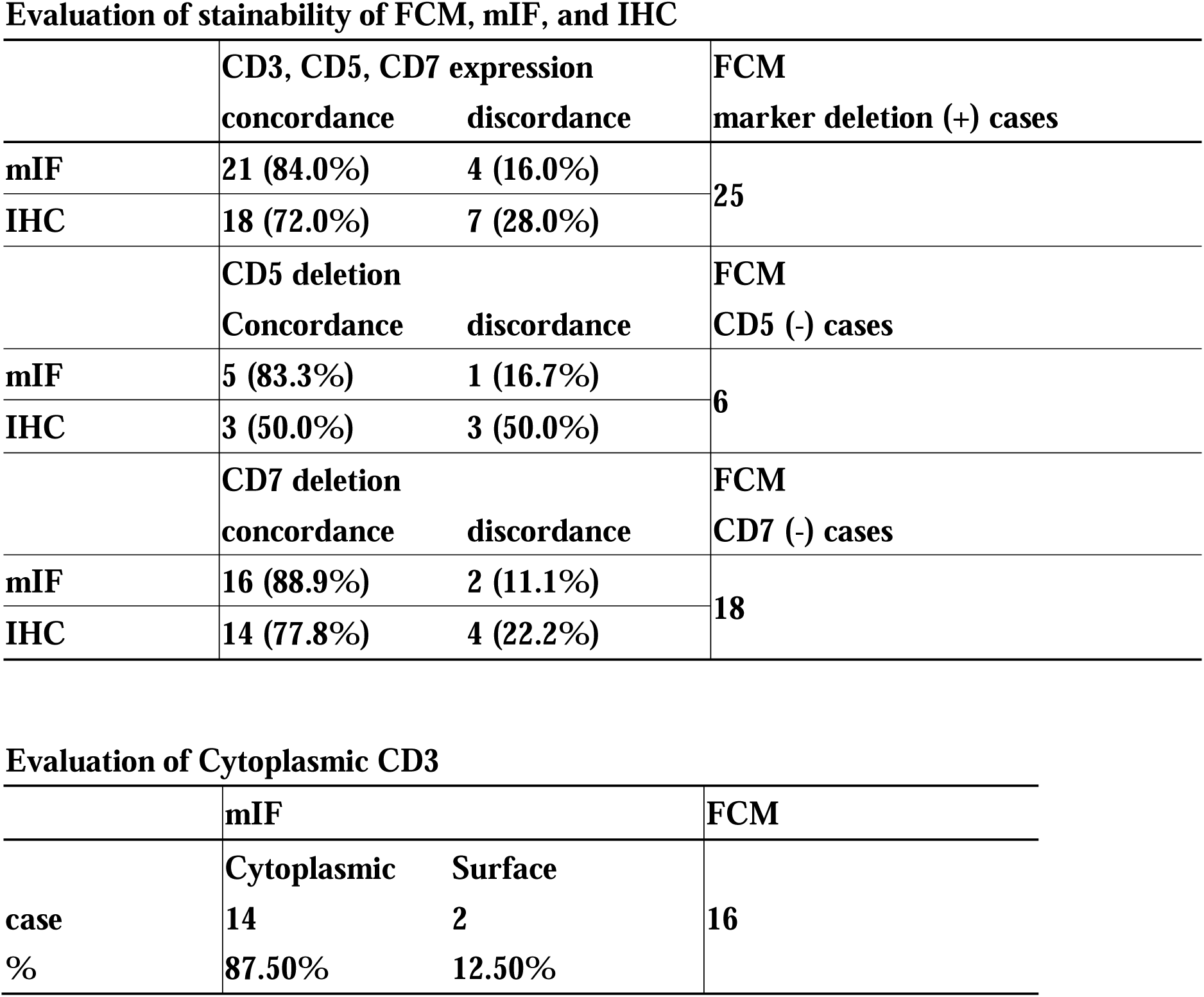
Evaluation of Stainability of FCM, multiplex immunofluorescence (mIF), and IHC.

CD3 staining in a target-like pattern inside the CD5 or CD7 membrane-positive images was observed, allowing visualization of cytoplasmic-CD3 (Fig. 5d). This finding was similar to the CD3 ζ intracytoplasmic expression in TCRα mutant T cells and was consistent with cytoplasmic-CD3 staining^29^. Of the 16 cases showing cytoplasmic-CD3 in FCM, similar staining was also confirmed in 14 cases (87.5%) in mIF (Table 1).

### jNMF-Based Spatial Clustering Analysis (jNMF-SCA)

The joint Non-Negative Matrix Factorization based Spatial Clustering Analysis (jNMF-SCA) was developed to categorize case samples and highlight characteristic regions within each classification to aid pathologists in phenotyping. jNMF, an algorithm for dimension reduction and feature extraction in multiplex analysis using multiple samples, facilitates multimodal analysis encompassing multidimensional data such as gene expression^30–34^. By decomposing multiple matrices into a common basis matrix W and feature values H, jNMF simplifies data complexity and reduces dimensionality, allowing the correlation pattern between multiple elements to be elucidated by projecting multiple matrices into a new shared space (Supplemental Fig. 5). To cluster samples by the spatial distribution of various phenotypic cells and to extract their characteristic regions using jNMF, whole slide images were segmented into subregions of 300 × 300 μm^2^, with the number of biomarker-positive cells quantified for each subregion. This study aimed to identify samples and regions containing cells with deletions of CD5 or CD7 in CD3-positive T-cell lymphoma. Subregions were organized by the total count of CD3 positive (+) cells, generating a matrix for the percentage of CD5 negative (-) cells or CD7 negative (-) cells, respectively, arranged as columns of sorted subregions and rows of samples (Fig. 6a). Subregions with a higher number of CD3(+) T cells were sequentially positioned from left to right in the matrix column. This ordering facilitated the interpretation of marker correlation, as the expression of CD3, CD5, and CD7 on normal T cells is correlated. Sorting the subregion space in descending order of CD3(+) cells highlighted the correlation among markers more straightforwardly.

**Figure 6.**
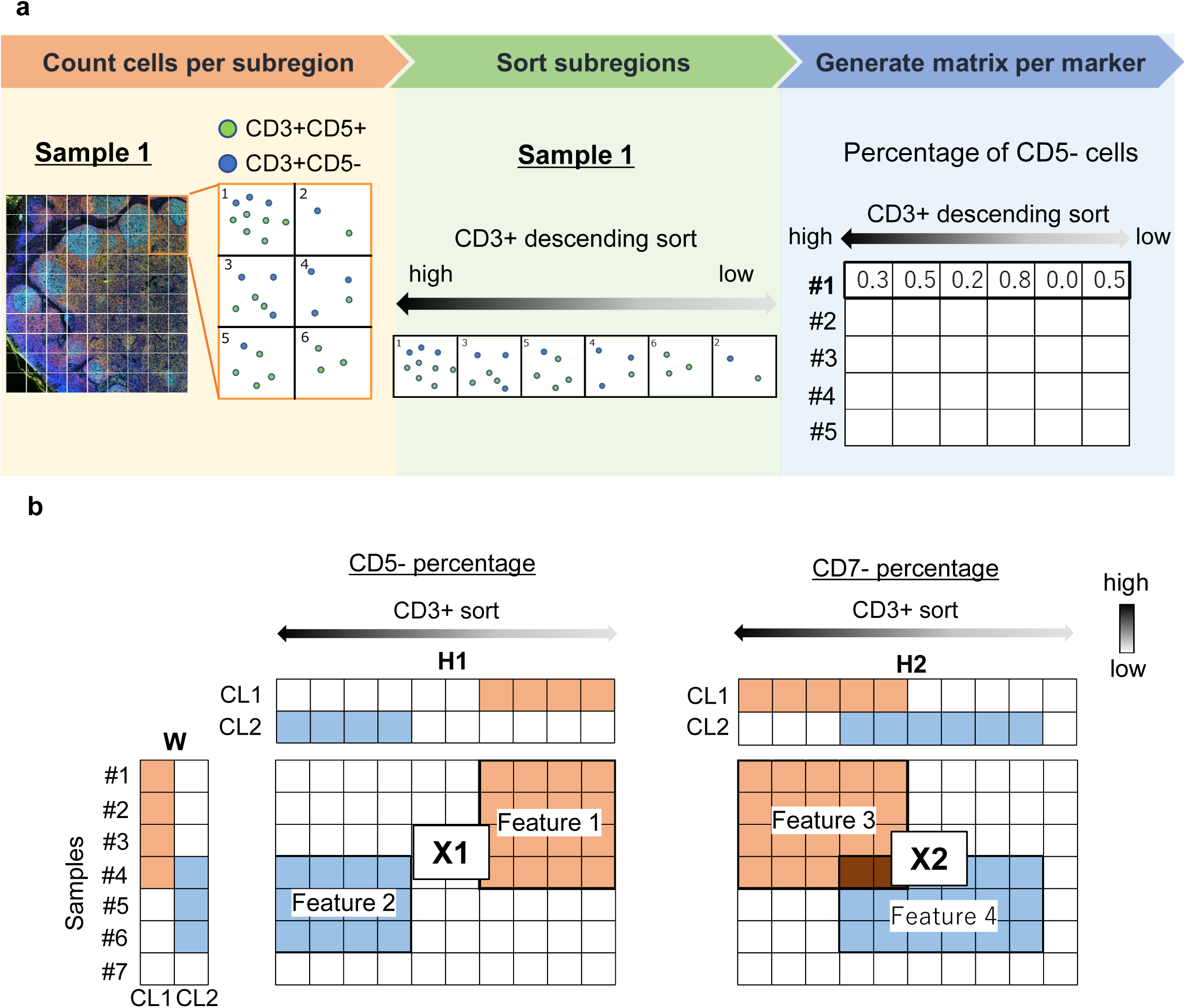
jNMF-Based Spatial Clustering Analysis (jNMF-SCA). (a) Data preprocessing flow for jNMF-SCA, detailing the division of the image into small regions of 300 × 300 µm^2^, counting positive cells for CD3, CD5, and CD7 in each region, and sorting regions by descending order of CD3(+) cell count. (b) Output image of jNMF-SCA, showing each matrix, X1 as a matrix of CD5 (-) ratio, and X2 as a CD7 (-) ratio. The first column of W corresponds to Cluster 1 (CL1), and the second column to Cluster 2 (CL2). Feature vectors of each cluster, CL1 and CL2, correspond to the first and second rows of H1 and H2, respectively. The orange and blue blocks represent characteristic regions of CL1 and CL2, respectively.

jNMF compresses the data’s dimensionality by decomposing several sample matrices into a common basis matrix W for clustering samples and a feature value H characterized by the distribution of multiple cell types (Fig. 6b). The jNMF algorithm uses the Euclidean distance and the updated equation employs the multiplicative update rule^30^. To illustrate the analysis results, X1 represents a matrix of the CD5(-) cell ratio sorted in descending order of the number of CD3(+) cells in each subregion, and X2 is a matrix of the CD7(-) cell ratio sorted similarly. The first column of W corresponds to Cluster 1 (CL1), indicating samples with high values of CD5(-) or CD7(-) cells in certain regions, and the second column corresponds to Cluster 2 (CL2). The feature vectors H for each cluster—CL1 and CL2—identify the regions with significant contributions for each cluster. The common factor matrix W aids in observing cluster classification, while the feature matrix H extracts regions with a high density of CD5(-) or CD7(-) cells, indicated by the orange and blue areas in the H matrix for CL1 and CL2, respectively (Fig. 6b).

### Assisted Diagnosis of T-cell Lymphoma Using jNMF-SCA

For the assisted diagnosis of T-cell lymphoma using jNMF-SCA, a total of 30 case samples, including 14 control samples and 16 T-cell lymphomas, were grouped into 2 clusters. Control samples were associated with low z-score assignments to either cluster, while T-cell lymphoma samples exhibited high z-score assignments to one or both clusters, as depicted in the W matrix. Cluster 1 (CL1) was characterized by CD5 deletions in regions with relatively few CD3(+) cells, as shown by H1, and CD7 deletions in regions with low to medium CD3(+) cell counts, as indicated by H2. Conversely, Cluster 2 (CL2) encompassed regions with consistent CD5 expression, as shown by H1, and a broad range of regions with CD7 deletions, as depicted by H2 (Fig. 7a).

**Figure 7.**
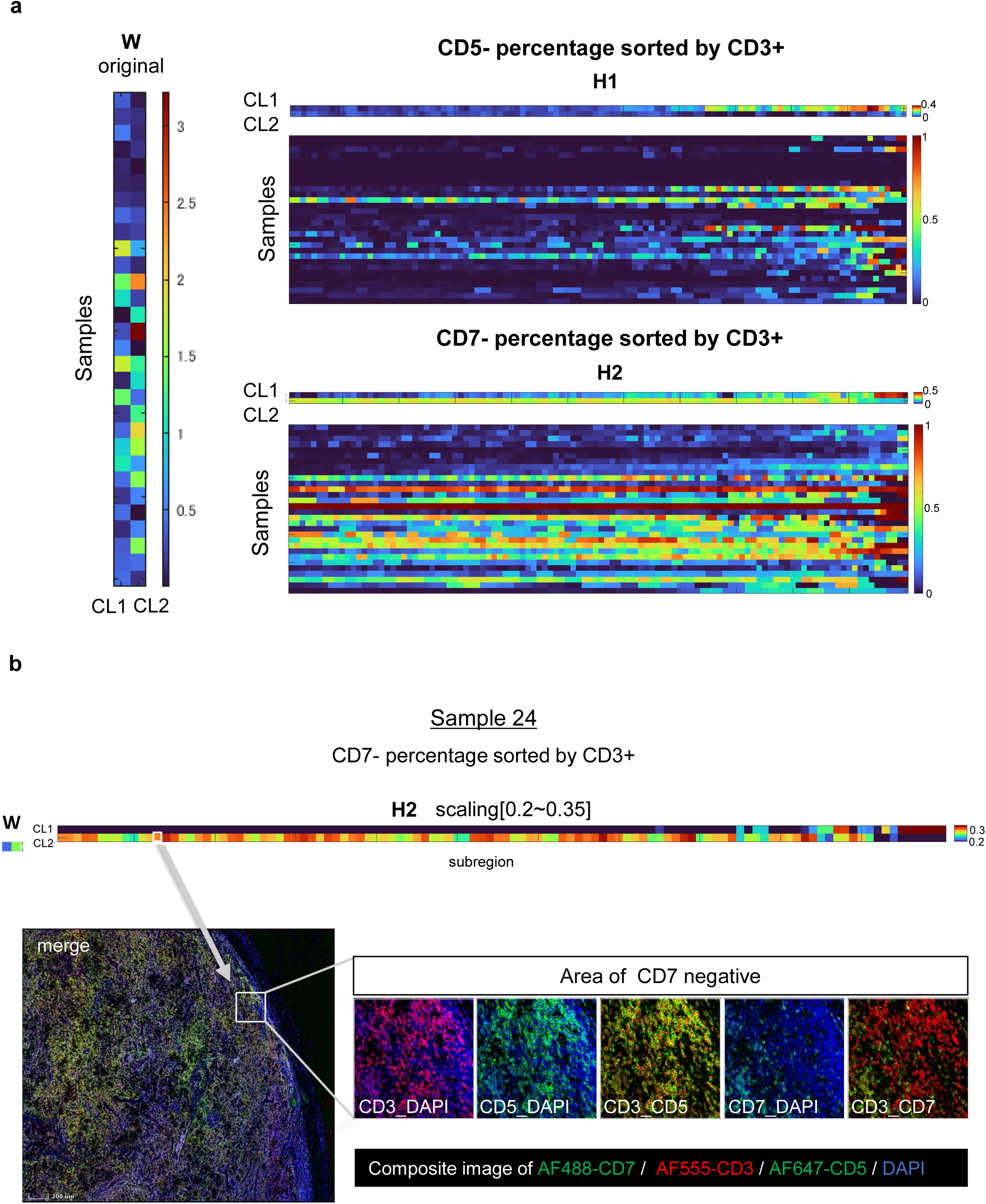
Assisted Diagnosis of T-cell Lymphoma using jNMF-SCA. (a) Dataset of lymphoma samples and results of jNMF-SCA (W, H), with a heatmap indicating higher values in red and lower values in blue. Regions with a high ratio of CD5 (-) or CD7 (-) appear in red. (b) Field of view with a high proportion of CD7 (-) indicated by jNMF (white box), showing merged and composite images of AF488-CD7 in green, AF555-CD3 in red, AF647-CD5 in green, and DAPI in blue.

To retrospectively detect samples suspected of harboring CD5 or CD7 negative T-cell lymphoma, a criterion was established defining such samples as those with a z-score in CL1 or CL2 of the W matrix exceeding the mean plus three times the standard deviation (μ+3σ) of the control sample. The threshold was calculated using control samples of the analyzed target, rather than using training data. This result was then compared to the subjective evaluation performed by the pathologist, who considered both dim detection and partial region negative detection as negative (Supplemental Table 3). Out of the 30 samples, including controls, 16 were identified by the pathologist as having CD5 or CD7 deletions, 15 of which were also detected by jNMF (true positives, TP). Furthermore, 14 specimens were deemed by the pathologist as triple positive, with 12 of these also classified as triple positive by jNMF (true negatives, TN). The analysis yielded a sensitivity of 93.75%, a specificity of 85.71%, and an overall correct response rate of 90.00% (Table 2). These outcomes demonstrate jNMF’s high sensitivity in detecting regions of deletion, including those considered dim or only affecting specific regions.

**Table 2.**
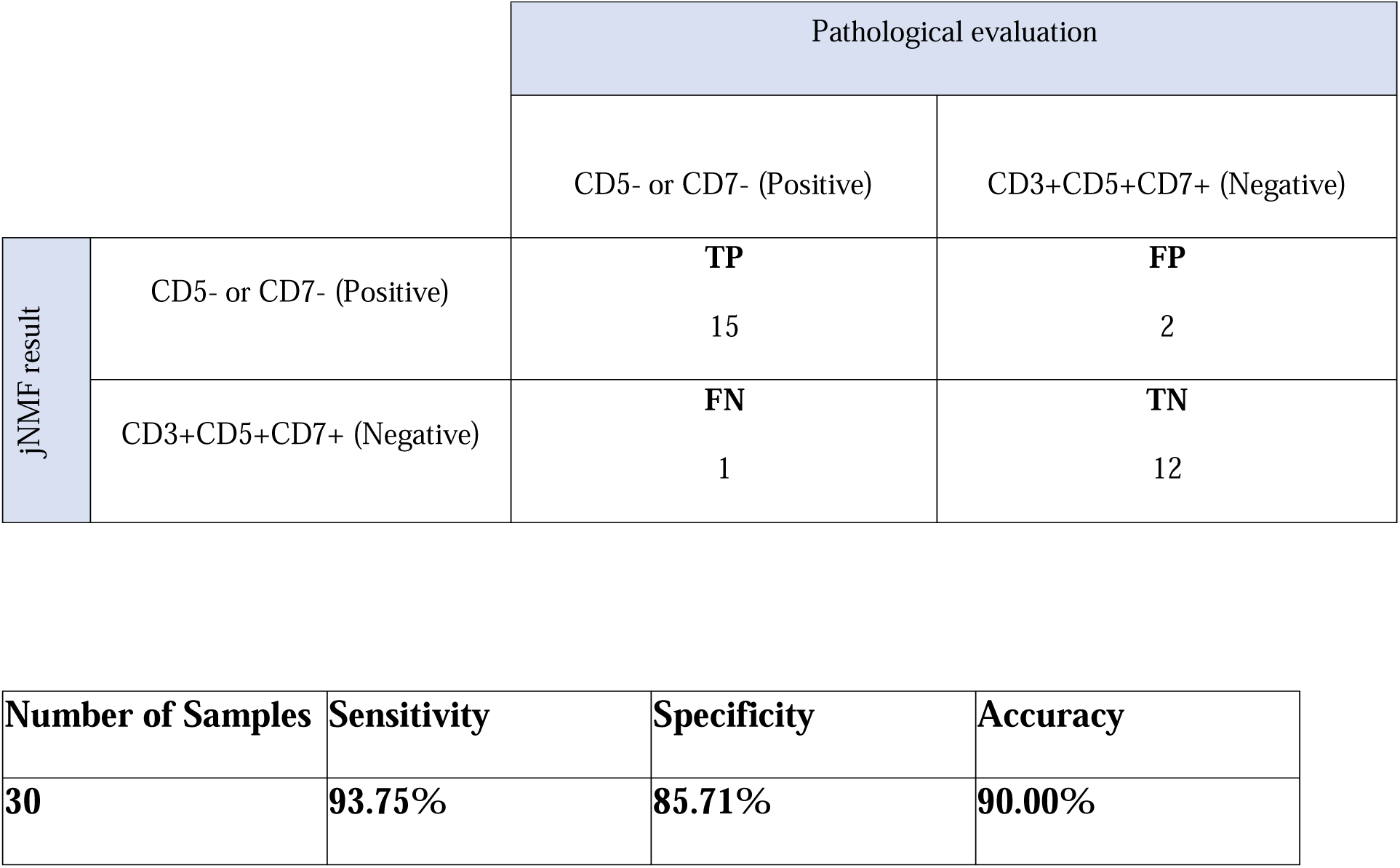
Correlation Between Pathologist Evaluation and jNMF Results. TP; True positive, FP; False positive, FN; False negative; and TN; True negative.

The pathologist’s phenotypic determinations were guided by the jNMF-SCA result display. For instance, Case sample 24 was assessed as CD3(+), CD5(+), CD7(-) by both the pathologists and jNMF. Focusing on the area with a high CD7(-) ratio as presented by jNMF-SCA facilitated the pathologist’s identification of the CD7 deletion area, which was not extensive across the entire field of view (highlighted by the white box in Fig. 7b). In the targeted field of view, this sample contained CD3-cytoplasmic positive cells that did not merge with either CD5 or CD7, but co-expression of CD5 was confirmed in the same cells, while numerous CD7 deletion cells were observed (Fig. 7b).

## Discussion

Here, we have developed spectral imaging acquisition and spectral separation technologies that differentiate fluorescent signals from autofluorescent signals, enabling multiple fluorescence imaging via the direct method. Accurate removal of autofluorescence is critical for the direct method and is ideally obtained from the specimen itself. A method for acquiring an autofluorescence spectrum by designating a region of interest (ROI) by an operator has also been proposed, however the acquired spectrum varies depending on the specified ROI ^35, 36^. SISS-mIF automatically acquires autofluorescence spectra, ensuring reproducibility. This approach allows simultaneous imaging of up to nine types of fluorescent markers, including 8 colors plus 4’,6-Diamidine-2’-phenylindole dihydrochloride (DAPI). The technology streamlines and accelerates data acquisition by enabling staining and scanning in single steps, simplifying the operational process, which is a crucial improvement for diagnostic workflows.

Moreover, akin to the capabilities of a flow-cytometer, our method’s multiplicity expands with the availability of primary antibodies tagged with fluorescent dyes, allowing staining and scanning in one step without complicating the operational process from start to finish or necessitating staining order consideration. Conversely, TSA and cyclic staining methods require additional steps for staining or imaging as the number of fluorescent colors increases, necessitating the optimization of staining order. This not only increases the time required for assay optimization and the entire assay process but also risks damaging the tissue section due to repeated solution reactions, potentially resulting in the loss of valuable specimens through peeling and defects. While similar one-step immunostaining has been proposed using microscope systems with bandpass fluorescence filters, limitations exist on the fluorophores that can be utilized to match the filter’s wavelength^37^. Our method, utilizing hyperspectral spectroscopy, NMF, and LSM, offers a broader selection of fluorophores due to its superior spectral separation capabilities. A notable challenge with one-step immunostaining is ensuring consistent antigen activation conditions, as some antibody clones necessitate specific activation conditions, like enzymatic reactions. Nevertheless, several antibodies require antigen activation under uniform conditions, akin to mass spectrometric tissue imaging. Consequently, SISS-mIF, capable of completing staining and imaging in a singular process, is optimally suited for diagnostic and laboratory workflows.

Our technique, which outputs images based on standardized intensity independent of capture conditions rather than fluorescent intensity, has demonstrated the capability to display images within a similar dynamic range, irrespective of image capture conditions or multiplex staining (Fig. 4). This method is particularly effective when combined with our spectral unmixing technique; otherwise, incomplete spectral unmixing, including autofluorescence, may lead to inaccurate standardized intensities. Anticipated to diminish batch effects in case sample analyses, this technique has indeed standardized data processing across all T-cell lymphoma case samples, regardless of acquisition times, including setting thresholds for marker expression. Photobleaching is another challenge when applying fluorescence imaging to diagnostic workflows. Unlike conventional fluorescence imaging or spectral imaging with wavelength tunable filter^12^, in which imaging is performed repeatedly by a band-pass filter set for each fluorescent molecule, this system minimized the effect of photobleaching due to uniform exposure time control across all fluorescent molecules, notwithstanding the disparate exposure times necessitated by repeated exposure and illumination for each fluorescent molecule.

The SISS-mIF shows potential for aiding lymphoma diagnosis, enabling precise co-expression evaluation of CD3, CD5, and CD7 within the same cell and facilitating accurate identification of biomarker expression locations, distinguishing between cell membrane and cytoplasm expressions (Fig. 6d). Moreover, it allows for the detection of regions with CD5 and CD7 expression deficiencies, offering analysis beyond phenotype distribution, akin to flow cytometry, by incorporating region information and potentially linking to malignant areas, such as those characterized by nuclear atypia. Overall, mIF was superior to IHC in all results, with a trend of IHC-negative cases being positive in mIF (Fig. 6). To enhance these outcomes, exploring cut-off values through standardized intensity with control samples may prove beneficial.

Additionally, our jNMF-SCA could notably support lymphoma diagnosis. Although high sensitivity is desirable, the analysis included one false negative (Table 2). Case sample 26, classified as a false negative, presented a challenging scenario as determined by the pathologist, exhibiting both full and partial positive expressions across CD3, CD5, and CD7. Notably, this sample exhibited higher standardized intensities for CD3, CD5, and CD7 than other case samples, falling below jNMF’s detection threshold using a common threshold (Supplemental Fig. 4).

Opting not to use jNMF-SCA for phenotype determination and instead employing its display output to guide the pathologist has also proven effective (Fig. 7b). This guidance functionality can efficiently and accurately pinpoint lesions, proving valuable for pathological evaluations, even in scenarios where a minimal number of lymphoma cells infiltrate the epidermis or adipose tissue in skin samples. Heatmaps generated by jNMF-SCA elucidate the negative rates of CD5 and CD7 in subregions, arranged in descending order of CD3 presence. Subregions towards the right tend to exhibit higher negative rates due to their peripheral tissue location, which naturally contains fewer T cells, leading to increased missing rates upon detecting autofluorescence (Fig. 7b). While there remains room for enhancement in data preprocessing for jNMF-SCA input and result output display, this methodology is anticipated to extend beyond T-cell lymphoma diagnosis assistance to other lymphomas and quantitative tumor microenvironment evaluation. Although synergizing with SISS-mIF improves positive/negative marker expression determination, jNMF-SCA could also complement other multiplex imaging methodologies. This adaptability allows jNMF-SCA not just to be paired with immunofluorescence but also to be applicable to hyperplex imaging, including spatial transcriptomics. Further studies are necessary to explore these applications^38–40^.

In summary, the combination of SISS-mIF and jNMF-SCA offers a robust framework for enhancing diagnostic accuracy and efficiency in lymphoma and potentially other cancers. SISS-mIF’s ability to accurately evaluate co-expression and precisely identify biomarker expression locations, coupled with jNMF-SCA’s high sensitivity and guidance functionality, presents a significant advancement in diagnostic methodologies. The utilization of heatmaps to display subregion-specific negative rates of CD5 and CD7 further aids in the nuanced analysis of lymphoma, allowing for a more detailed examination of the tumor microenvironment. While both SISS-mIF and jNMF-SCA show great promise individually, their combined use could revolutionize the approach to lymphoma diagnosis, providing pathologists with a powerful toolset for identifying and characterizing disease with unprecedented precision.

## RESOURCE AVAILABILITY

### Lead contact

Requests for further information and resources should be directed to and will be fulfilled by the lead contact, Kazuhiro Nakagawa.

### Materials availability

All unique/stable reagents generated in this study are available from the lead contact without restriction.

### Data and code availability

The data that supports the findings of this study are available within the article and its Supplementary Materials. The raw image data will be shared on reasonable request to the corresponding author.

## Acknowledgements

We thank Shuichi Kakuda and Chinatsu Ito for their contribution to staining slides and capturing imaging data.

## Author contributions

T.N., N.K., and T.T. contributed equally to this work. T.N. and K.N. designed and conceived this study. T.K., H.T., developed spectral imaging system. T.N. performed experiments and collected data. T.N., N.K., K.I. M.S. analyzed data. M.K. designed clinical study. T.T. M.I. I.O. M.K. provided clinical samples and clinical information. T.T. I.O. M.K. conducted pathological evaluation and validation. T.N. N.K. T.T. K.I. T.K. M.K. K.N. wrote the manuscript. K.N. supervised the project. All authors have read and agreed to the published version of the manuscript.

## Methods

### Tissue Slides

For multiplex immunofluorescent staining, we utilized human tonsil tissue, human skin tissue, and human breast tissues, which were diagnosed as reactive follicular hyperplasia, nodular melanoma, and infiltrating ductal carcinoma, respectively. These tissue specimens were acquired from BioIVT, LLC (Westbury, NY), with informed consent obtained from the source, excluding personal information. For the T-cell lymphoma study, twenty-five FFPE specimens from patients with T-cell lymphoma at Tokyo Medical and Dental University Hospital (Tokyo, Japan), spanning from 2014 to 2021 with accompanying flow cytometry data, were examined. Control samples comprised 10 non-neoplastic lymph node specimens and 3 inflamed skin specimens from the same institution. This study received approval from the institutional ethics board (M2019-204). All tissues were sectioned at 4-μm thickness and placed on CREST slide glass (Matsunami Glass Ind., Ltd., Osaka, Japan).

### Immunostaining

IF and IHC assays were conducted using a fully automated stainer, BOND RXm (Leica Biosystems, Nussloch, Germany), according to the manufacturer’s guidelines. Slides underwent de-paraffinization with Bond Dewax Solution (Leica Biosystems) for 30 min at 60 °C. Epitope retrieval was performed using BOND Epitope Retrieval Solution 2 (Leica Biosystems) for 20 min at 100 °C. Primary antibodies, as listed in Supplemental Table 4, were diluted with BOND Primary Antibody Dilution (Leica Biosystems) to a specific concentration and applied to the slides, which were then incubated at room temperature for 30 min. For brightfield detection, slides were stained with DAB and hematoxylin utilizing BOND Polymer Refine Detection (Leica Biosystems) as per the manufacturer’s instructions, followed by dehydration, permeabilization, and mounting with Pathomount (FUJIFILM Wako Pure Chemical Corp., Osaka, Japan) and a cover glass. For fluorescent detection, slides underwent antibody incubation followed by DAPI staining and were mounted with ProLong Diamond Antifade Mountant (Thermo Fisher Scientific, Waltham, MA, USA) and a cover glass.

### Hyperspectral Imaging

The line-scan spectrum imaging system was assembled from a commercially available BX63 microscope (Olympus, Tokyo, Japan) with a x20 objective lens, a laser light source, and a custom-made monochromator. Four lasers were utilized for excitation (405 nm: NDV7116(Nichia corp., Tokushima, Japan), 488 nm: NDS7175E (Nichia corp.), 532 nm: VFL-P-1000-532-OEM1-B1(MPB Communications Inc.), 638 nm: HL63193MG (Ushio Inc., Tokyo, Japan). An emitted laser was shaped into a flat-top beam profile using a cylindrical lens array, a scanning mirror to negate any interference pattern, and a custom-made monochromator (Supplemental Fig. 1a). An Offner-type monochromator was selected for imaging spectroscopy, casting spatial and wavelength axes in two dimensions on an image sensor in a single exposure with a custom grating (Supplemental Fig. 1b). The wavelength resolution of the Offner-type monochromator was set at 8 nm. A CMOS camera captured the spectrum image. Scans were conducted with each of the 4-excitation lasers. An excitation filter was selected for each laser (ZET405/20x, ZET488/10x, ZET532/10x, ZET642/20x (Chroma Technology corp., Bellows Falls, VT). Dichroic mirrors and emission filters were chosen with a 4-band filter respectively (ZT405/488/532/640rpc-XT, ZET405/488/532/642m (Chroma Technology corp.).

### Reference Spectrum Generation

Reference spectra were obtained by imaging standard samples encapsulated with fluorescent antibodies with a hyperspectral imaging system. Standard samples were prepared by uniformly dispersing the fluorescent antibody in the mounting medium, applying it to a glass slide, and mounting the coverslip to a thickness of 20 μm. The reference spectrum was normalized to the intensity per molecule/voxel from the concentration of the fluorescent antibody dispersed in the mounting medium and the voxel volume corresponding to one pixel of the imaging.

### Spectral Unmixing

Unmixing was performed in the order described in Algorithm 1 as follows: Step 1: Open a file in an original file format using MFICo_dat_converter function.

Steps 2–5: Since the data is output for each unit size, the Gram matrix is calculated and added by turning the for loop for the number of data items.

Step 6: The tAA_sum is decomposed by the least squares method (LSM) using the initial spectrum (W_0) to obtain the coefficient (H_0).

The initial spectrum (W_0) is the spectrum (Arachidonic Acid, Elastin, FAD, PLD, Protoporphyrin) considered as autofluorescence.

Step 7: The initial spectrum (W_0) and its coefficient (H_0) are decomposed by the nonnegative matrix factorization (NMF) to obtain the optimized autofluorescence spectrum (W).

Steps 8–9: The obtained optimized autofluorescence spectra and the reference spectra of fluorescent antibodies used for staining are decomposed by the least squares method (LSM) in the for loop of the data number to obtain the unmixing result.

Step 10: The result obtained in each loop is saved as a tif image by the save_tif function.

### Algorithm 1

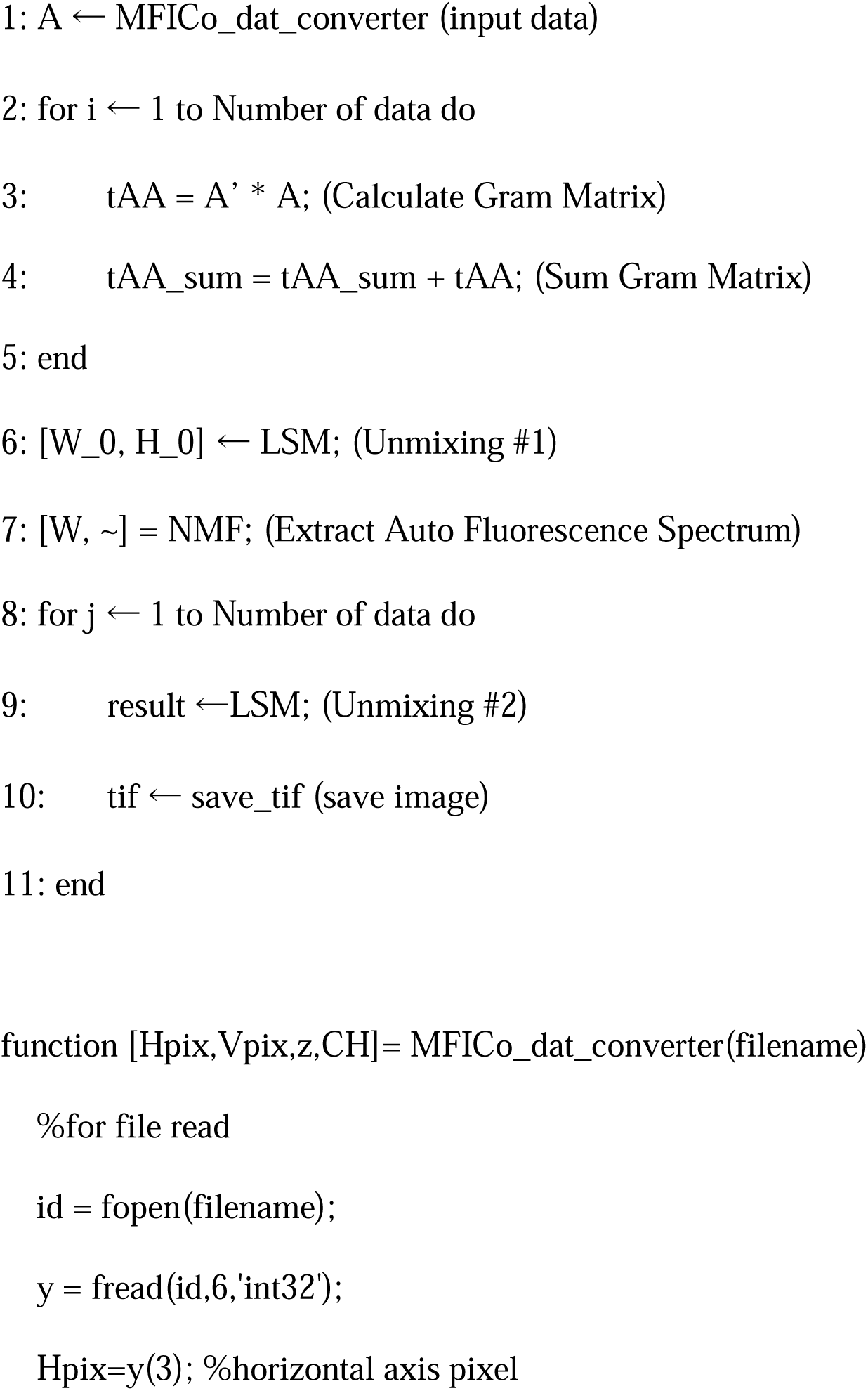
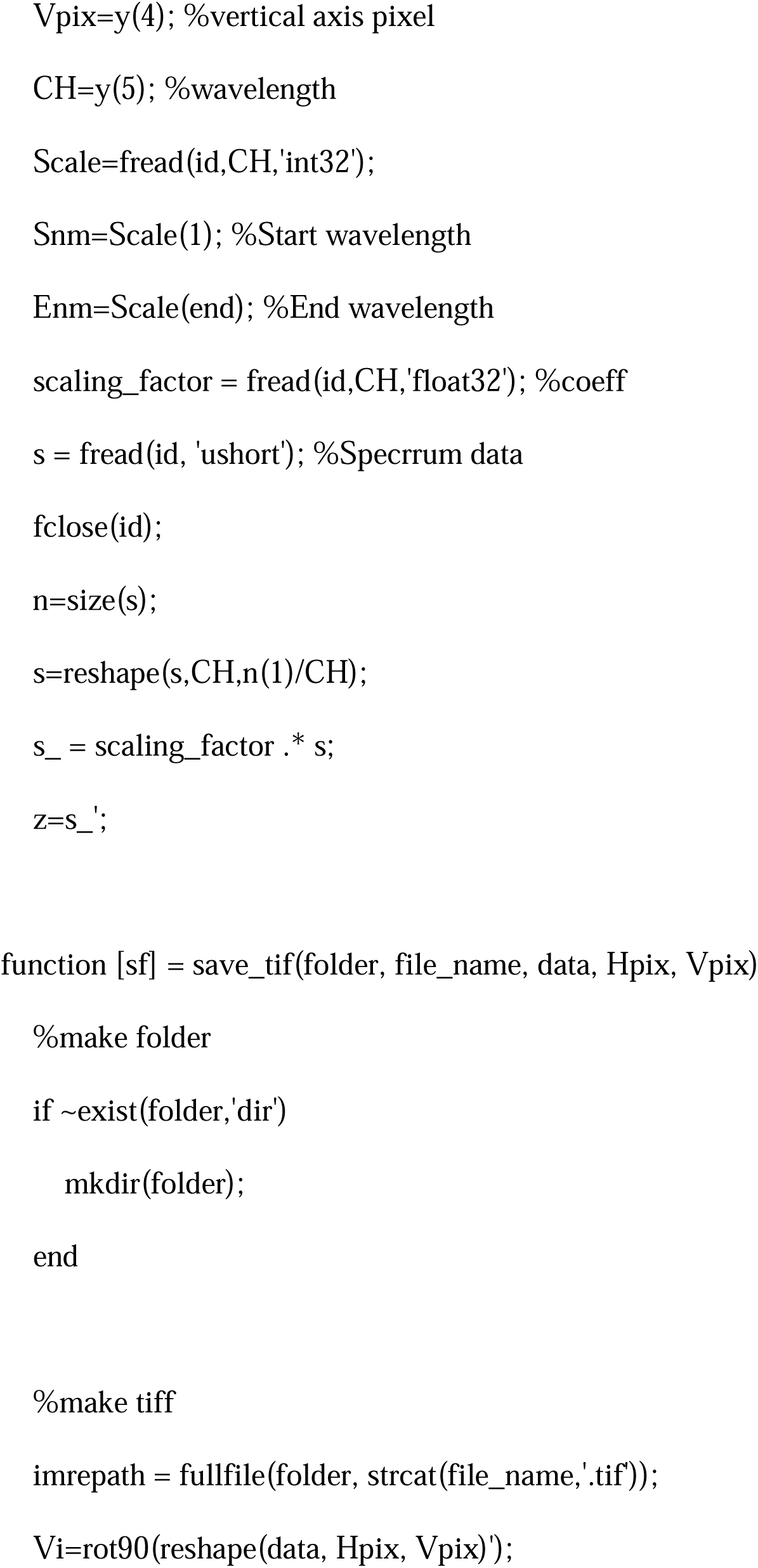
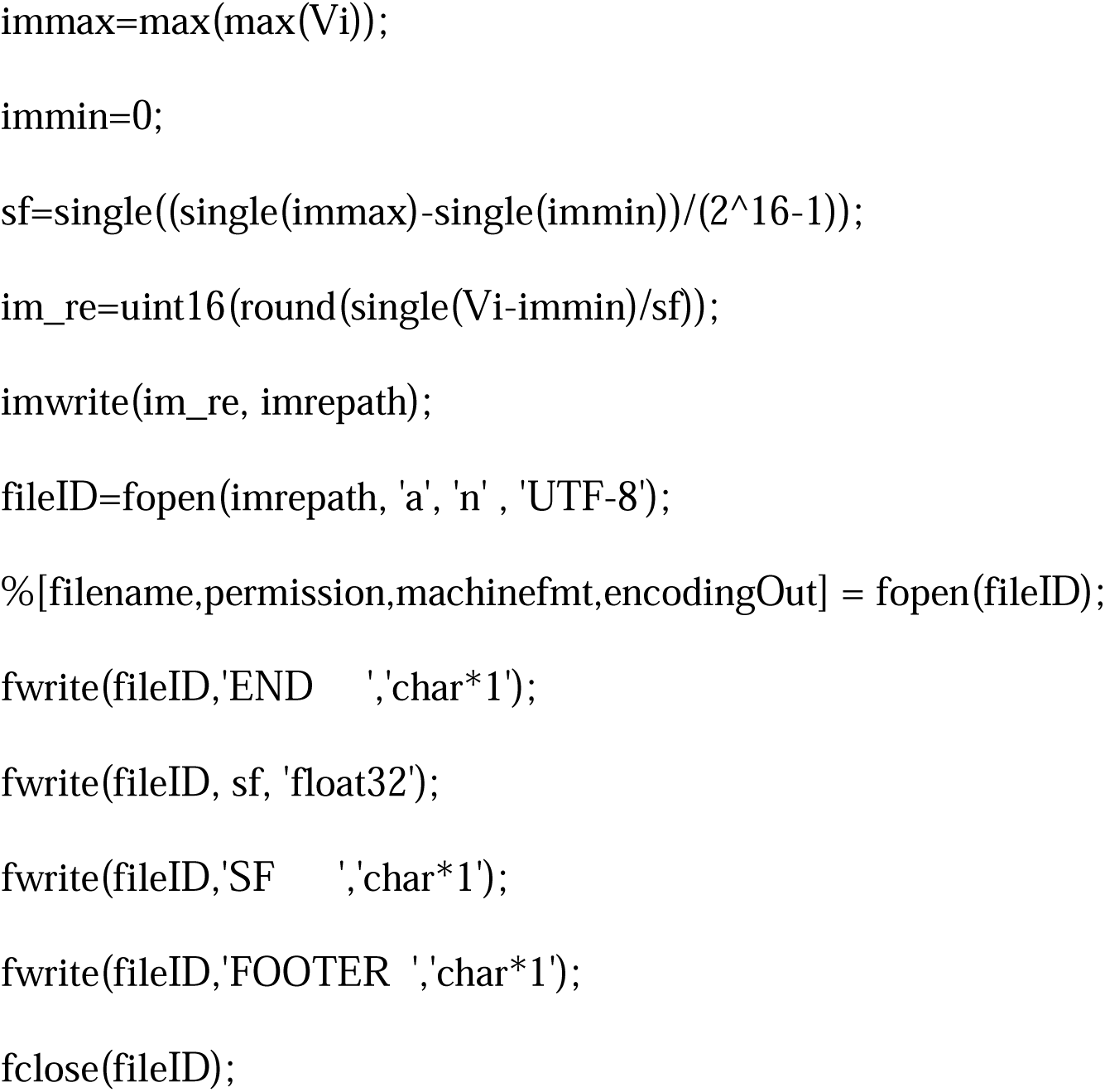

### Image Shading Correction

Shading correction was conducted by deriving the shading pattern from all spectral cubes acquired by the system and incorporating the reciprocal values into each spectral cube. Following shading correction, spectral unmixing was performed on the corrected spectrum cube using the reference spectrum.

### Image Deconvolution

A 1-μm pinhole light source with a defined structure was imaged using a line-scan spectral imaging system to capture its intensity distribution. The device’s intrinsic point spread function was identified by fitting the ideal intensity distribution of a 1-μm aperture into the actual intensity distribution captured by the device. Subsequently, the image underwent processing with Richardson-Lucy deconvolution^41, 42^, using the derived point spread function.

### Image Stitching

Unmixing occurred for each default size (2432 × 640 pix), and stitching was employed for a comprehensive field of view. Each default size data included a gloss of approximately 5% (128 pix) per side, and deviations in the x and y directions were calculated by template matching within that region^43^. A wide-field image was compiled by accounting for these deviations and performing stitching.

### Image Cytometry Analysis

Stitched images were segmented into small regions (blocks) of 300 × 300 µm^2^. Following cell segmentation and expansion treatments, the number of antibodies per cell for biomarkers (CD3, CD5, and CD7) was calculated. Cell segmentation utilized nuclear detection algorithm based on DAPI signal, and the expansion process extended 4 pixels from the nucleus. The standardized intensity independent of capture conditions was ascertained by averaging pixels corresponding to the expanded area. Cells exceeding the antibody number threshold for each biomarker were identified as positive cells. The threshold for determining positive cells for each biomarker was set at the 98^th^ percentile of antibody numbers per DAPI-positive cell in the unmixed images assigned to each marker in non-immunostained images of control samples. This approach to determining positive cells by standardized intensity, set at thresholds for each marker, resulted in more than 85% of CD3 positive cells being concurrently positive for CD3, CD5, and CD7 in immunostained images of all control samples, underscoring the method’s precision.

### Data Imputation for jNMF-SCA

To construct the matrix for jNMF-SCA, aligning the number of rows (i.e., the number of subregions) is essential. jNMF’s effectiveness in clustering can be influenced when more than 10% of the data is completed with zeros^33^. In this study, the number of subregions for all case samples was standardized to 110, the maximum value among the samples, for all 30 case samples. The imputed row data were augmented by randomly extracting subregions until reaching a total of 110, ensuring uniformity across samples.

### Extracting Cluster Membership After jNMF-SCA

Following jNMF, the resulting matrix W was standardized using Z-scores to facilitate comparison and analysis. The Z-scores of matrix W were calculated using the following equation:

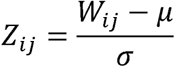

where µ represents the mean value of W, and σ represents the standard deviation of W. i represents the row number and j represents the column number. This standardization allowed for the evaluation of cluster membership based on statistical significance. The population of Z-scores for control samples (Sample #1∼Sample #9) for each cluster was estimated using the maximum likelihood method, from which the mean value µ and standard deviation σ of the estimated population were determined. A cutoff value of µ+3σ was then set for each cluster, with samples exceeding this value classified within clusters and considered as negative judgments in jNMF, enhancing the method’s specificity in cluster determination.

### Phenotyping by Pathologist

Phenotype determination using fluorescence imaging was performed by three pathologists employing an in-house image viewer, setting the threshold for each marker at the 99.8^th^ percentile of the standardized intensity in the fluorescent images, excluding autofluorescence components. Two rounds of evaluation were performed, leveraging the guidance function of jNMF-SCA. Discrepancies between the first and second round evaluations prompted a discussion among the pathologists, culminating in a consensus decision. In comparing results with jNMF, evaluations by the pathologists were aggregated as negative even if detected partially negative or classified as dim, aiming for a conservative and accurate diagnostic approach. This methodological rigor ensures a high level of precision in lymphoma diagnosis, facilitating effective treatment planning and patient care.

### Generation of Superimposed Simulation Images

Superimposed simulation images were generated using the following procedure.

#### 1 Determination of the intensity of the superimposed fluorescent spectrum image

The intensity of the fluorescent spectra to be superimposed to the background image (an unstained image with only DAPI staining) was determined based on the intensity of standard samples and of DAPI-stained tonsil tissue slide.

i. Calculation of the peak position intensity of fluorescence: The 16 nm wide intensity around the peak position of each fluorescent spectrum was obtained by summing the intensities of the peak 2 channels of the standard sample.
ii. Calculation of peak position intensity of autofluorescence: The 16 nm wide intensity of the autofluorescence in the background image was obtained by summing the intensities of the 2 channels corresponding to each fluorescent spectrum used in (i).
iii. Determination of the fluorescent intensity to be superimposed to the background image: The multiplication factor n, which was obtained by the fluorescence intensity in (i), was twice that of the autofluorescence spectrum obtained in (ii), was determined.

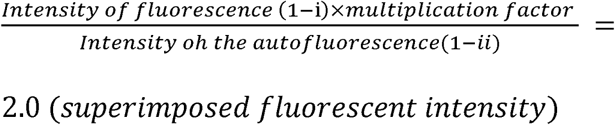

Equation 1: Determining the fluorescent intensity superimposed to background image

#### 2 Generation of simulation image

Nine standard samples (AF488-CD4, AF532-CD5, AF555-CD3, AF568-Ki67, eF615-CD20, AF647-CD8, AF700-Bcl2, AF750-CD68, AF790-CD7) were used to generate the simulation images. Tile image of the spectrum cube to which shot noise was applied were generated and superimposed on the background image according to the following procedure.

i. Adjust the intensity of the fluorescent spectrum relative to the autofluorescence intensity of the background image by multiplying the multiplication factor obtained in (1-iii).
ii. Convert spectral radiance to AD value by dividing the wavelength calibration data for conversion to spectral radiance.
iii. Convert AD values to charge quantity (e-) based on the gain at the time of background image capture and the saturation capacity of the pixels. 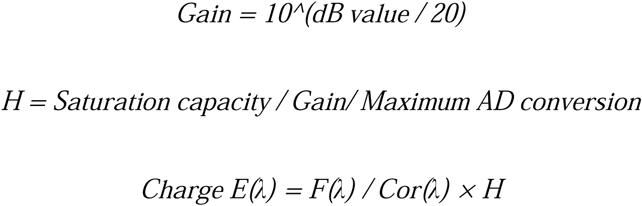 Equation: Charge quantity conversion equation F(λ): Standard spectrum of the fluorescence Cor(λ): Wavelength calibration data H: Conversion coefficient E(λ): Charge quantity
iv. Apply random noise σ = √S (S: charge quantity per pixel in electrons) as shot noise 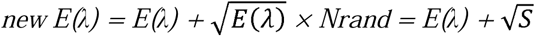 newE(λ): Standard spectrum of the fluorescence overlaid with shot noise Nrand: Normally distributed random number with σ=1 S: Charge quantity per pixel in electrons
v. Convert the charge quantity (e-) back to spectral radiance using the reverse process of (i)∼(iii)
vi. Generate a 10x10 pixel spectral cube with 1 pixel of spectral radiance generated in (v) for each fluorescence.
vii. Generate an image in which multiple spectral cubes generated in (vi) are placed on stripes.
viii. Superimpose the striped image generated in (vii) to the background image.

