## Supplemental text for "Spatial Clustering Analysis with Spectral Imaging-based Single-Step Multiplex Immunofluorescence (SISS-mIF) for Assisting Histological Diagnosis"

**Supplemental Table 1. Diagnostic findings for general T-cell lymphoma.**

| Methods | Details |
| --- | --- |
| Morphological observation | Atypia of lymphoid cells  Loss of structure of lymph nodes |
| Immunohistochemical analysis | Lineage decision (CD3-positive),  Aberrant expression or deletion |
| Flowcytometry | Deletion of common T-cell surface markers,  i.e., CD3, CD5, CD7 |
| Genetic analysis | T-cell receptor rearrangements |

**Supplemental Table 2. Comparison results of FCM, IHC, and mIF.**

|  | **Position** | **Diagnosis** | **FCM** | **m-IF** | **FCM/m-IF** | **IHC** | **FCM/IHC** | **Cytoplasmic CD3** | |
| --- | --- | --- | --- | --- | --- | --- | --- | --- | --- |
|  |  |  |  |  |  |  |  | **FCM** | **m-IF** |
| case 1 | lymph node | AITL | CD5(+)CD7(-) | CD5(+)CD7(-) |  | CD5(+)CD7(-) |  | + | + |
| case 2 | skin | C-TCL | CD5(+)CD7(-) | CD5(+)CD7(-) |  | CD5(+)CD7(-) |  |  | + |
| case 3 | lymph node | PTCL-NOS | CD5(+)CD7(+) | CD5(+)CD7(+) |  | CD5(+)CD7(+) |  | + | + |
| case 4 | lymph node | PTCL | CD5(-)CD7(-) | CD5(-)CD7(-) |  | CD5(+)CD7(+)/CD5(-)CD7(-) | mismatch |  |  |
| case 5 | skin | C-TCL | CD5(+)CD7(-) | CD5(+)CD7(-) |  | CD5(+)CD7(-) |  | + | + |
| case 6 | skin | PTCL-NOS | CD5(+)CD7(-) | CD5(+)CD7(-) |  | CD5(+)CD7(-) |  |  |  |
| case 7 | skin | C-TCL | CD5(+)CD7(-) | CD5(+)CD7(-) |  | CD5(+)CD7(-) |  | + | + |
| case 8 | skin | Extranodal NK/T CL | CD5(-)CD7(+) | CD5(-)CD7(-) | mismatch | CD5(-)CD7(-)/CD5(-)CD7(+) | mismatch |  |  |
| case 9 | lymph node | PTCL-NOS | CD5(+)CD7(-) | CD5(+)CD7(-) |  | CD5(+)CD7(-)/CD5(+)CD7(+) | mismatch |  | + |
| case 10 | tonsil | C-TCL | CD5(-)CD7(+) | CD5(-)CD7(+) |  | CD5(+)CD7(+)/CD5(-)CD7(+) | mismatch | + | + |
| case 11 | lymph node | C-TCL | CD5(+)CD7(-) | CD5(+)CD7(-) |  | CD5(+)CD7(-) |  | + | + |
| case 12 | skin | PTCL-NOS | CD5(+)CD7(-) | CD5(+)CD7(-) |  | CD5(+)CD7(-) |  | + | + |
| case 13 | skin | PTCL-NOS | CD5(-)CD7(-) | CD5(-)CD7(-) |  | CD5(-)CD7(+)/CD5(-)CD7(-) | mismatch |  | suspected |
| case 14 | lymph node | C-TCL | CD5(+)CD7(-) | CD5(+)CD7(-) |  | CD5(+)CD7(-) |  | + | + |
| case 15 | skin | AITL | CD5(-)CD7(+) | CD5(-)CD7(+) |  | CD5(-)CD7(+) |  | + | + |
| case 16 | lymph node | PTCL | CD5(+)CD7(-) | CD5(+)CD7(-) |  | CD5(+)CD7(-)/CD5(+)CD7(+) | mismatch |  |  |
| case 17 | skin | AITL | CD5(+)CD7(-) | CD5(+)CD7(-) |  | CD5(+)CD7(-) |  | + | + |
| case 18 | skin | PTCL-NOS+B-LPD | CD5(-)CD7(+) | CD5(+)CD7(+) | mismatch | CD5(+)CD7(-)/CD5(-)CD7(+) | mismatch | + |  |
| case 19 | skin | AITL | CD5(+)CD7(-) | CD5(+)CD7(-) |  | CD5(+)CD7(-) |  |  |  |
| case 20 | lymph node | TCL | CD5(+)CD7(-) | CD5(+)CD7(-) |  | CD5(+)CD7(-) |  |  | + |
| case 21 | lymph node | TCL | CD5(+)CD7(-) | CD5(+)CD7(+) | mismatch | CD5(+)CD7(-) |  |  | + |
| case 22 | lymph node | TCL and classical Hodgkin Lymphoma | CD5(+)CD7(+) | CD5(+)CD7(+) |  | CD5(+)CD7(+) |  | + |  |
| case 23 | lymph node | TCL | CD5(+)CD7(-) | CD5(+)CD7(+) | mismatch | CD5(+)CD7(-) |  |  | + |
| case 24 | lymph node | TCL | CD5(+)CD7(+) | CD5(+)CD7(+) |  | CD5(+)CD7(+) |  | + | + |
| case 25 | lymph node | TCL | CD5(+)CD7(-) | CD5(+)CD7(-) |  | CD5(+)CD7(-) |  |  |  |
| **AITL, Angioimmunoblastic T-cell lymphoma; TCL, T-cell lymphoma; C-TCL, Cutaneous T-cell lymphoma; PTCL, Peripheral T-cell lymphoma; NOS, Not otherwise special** | | | | | | |  |  |  |

**Supplemental Table 3. Comparative results between jNMF and pathological evaluation.**

| **Sample** | **jNMF (W_Zscore)** | | **Fluorescence evaluation by pathologists** |
| --- | --- | --- | --- |
|  | **CL1** | **CL2** |  |
| Sample 1 | -0.408711 | -0.837104 | CD3+CD5+CD7+ |
| Sample 2 | -0.613431 | -0.708604 | CD3+CD5+CD7+ |
| Sample 3 | -0.193851 | -0.746938 | CD3+CD5+CD7+ |
| Sample 4 | -0.872739 | -0.700534 | CD3+CD5+CD7+ |
| Sample 5 | -0.75731 | -0.910162 | CD3+CD5+CD7+ |
| Sample 6 | -0.786066 | -0.800953 | CD3+CD5+CD7+ |
| Sample 7 | -0.673803 | -0.73592 | CD3+CD5+CD7+ |
| Sample 8 | -0.339582 | -0.515589 | CD3+CD5+CD7+ |
| Sample 9 | -0.672172 | -0.606108 | CD3+CD5+CD7+ |
| Sample 10 | 1.78626 | 0.148457 | CD3+CD5+CD7- |
| Sample 11 | -0.455719 | -0.747316 | CD3+CD5+CD7+ |
| Sample 12 | 1.181525 | 2.577072 | CD3+CD5-CD7- |
| Sample 13 | 0.589406 | -0.580529 | CD3+CD5+CD7- |
| Sample 14 | -0.910162 | 0.729312 | PartialyCD3+CD5+CD7- |
| Sample 15 | -0.516434 | 3.836946 | CD3+CD5+CD7- |
| Sample 16 | -0.121595 | -0.910162 | CD3+CD5+CD7+ |
| Sample 17 | 1.717482 | 0.87195 | CD3dimCD5dimCD7dim |
| Sample 18 | -0.762072 | 0.590998 | CD3+CD5+CD7- |
| Sample 19 | 1.129783 | -0.297946 | CD3+CD5+CD7+ |
| Sample 20 | -0.63902 | 0.954227 | CD3+CD5-CD7- |
| Sample 21 | -0.347789 | 2.009059 | CD3+CD5dimCD7dim |
| Sample 22 | 0.433956 | 1.47826 | CD3＋CD5-CD7dim |
| Sample 23 | 0.711321 | 0.061656 | CD3+CD5+CD7+ |
| Sample 24 | -0.299853 | 1.28717 | CD3+CD5+CD7- |
| Sample 25 | -0.660011 | -0.107375 | CD3+CD5dimCD7dim |
| Sample 26 | -0.287476 | -0.910162 | CD3+CD5+CD7+, CD3+CD5dimCD7dim |
| Sample 27 | -0.733545 | -0.142187 | CD3+CD5dimCD7dim |
| Sample 28 | -0.4243 | 1.285274 | CD3+CD5+CD7dim |
| Sample 29 | -0.357542 | -0.294256 | CD3+CD5dimCD7- |
| Sample 30 | -0.289958 | -0.705129 | CD3+CD5+CD7+ |

*Cells exceeding the cutoff value are colored blue.

**Supplemental Table 4. List of primary antibodies used.**

| Figure | Panel | Dye | Target | Locations | Clone | Concentration μg/ml | Species | Manufacturer | Catalog number |  |
| --- | --- | --- | --- | --- | --- | --- | --- | --- | --- | --- |
| Fig. 4a | first | AF488 | CD4 | membrane | EPR6855 | 5 | Rabbit | Abcam | ab196372 |  |
|  |  | AF555 | PD-1 | membrane | NAT-105 | 5 | Mouse | Abcam | ab280864 |  |
|  |  | AF568 | Ki-67 | nucleus | EPR3610 | 1 | Rabbit | Abcam | ab211968 |  |
|  |  | AF647 | PD-L1 | membrane | 73-10 | 5 | Rabbit | Abcam | ab237403 |  |
|  |  | AF680 | CD3 | membrane | EP4426 | 5 | Rabbit | Abcam | ab226073 | conjugated using labeling kit |
|  |  | AF700 | Cytokeratin | cytoplasm | AE1-AE3 | 1 | Mouse | Novus Biologicals | NBP2-33200AF700 |  |
|  |  | AF750 | CD8 | membrane | C8-144B | 5 | Mouse | Cell Signaling Technology | 53912BC |  |
|  |  | AF790 | CD68 | cytoplasm | KP-1 | 5 | Mouse | Santa Cruz Biotechnology | sc-20060-AF790 |  |
| Fig. 4b | second | AF488 | CD45RO | membrane | UCHL1 | 5 | Mouse | BioLegend | 304212 |  |
|  |  | AF555 | CD3 | membrane | EP4426 | 1 | Rabbit | Abcam | ab208514 |  |
|  |  | AF647 | PD-1 | membrane | NAT-105 | 5 | Mouse | Abcam | ab220301 |  |
|  |  | AF680 | SOX10 | nucleus | SP267 | 1 | Rabbit | Abcam | ab245760 | conjugated using labeling kit |
|  |  | AF750 | Cytokeratin | cytoplasm | AE1-AE3 | 5 | Mouse | Novus | NBP2-33200AF750 |  |
| Fig. 4c | third | AF488 | CD4 | membrane | EPR6855 | 5 | Rabbit | Abcam | ab196372 |  |
|  |  | eF570 | Foxp3 | nucleus | 236A-7E | 5 | Mouse | Thermo | 41-4777-82 |  |
|  |  | eF615 | CD20 | membrane | L26 | 1 | Mouse | Thermo | 42-0202-82 |  |
|  |  | AF647 | CD8 | membrane | C8-144B | 5 | Mouse | BioLegend | 372906 |  |
|  |  | AF680 | CD68 | cytoplasm | KP-1 | 5 | Mouse | Santa Cruz Biotechnology | sc-20060-AF680 |  |
|  |  | AF750 | Cytokeratin | cytoplasm | AE1-AE3 | 5 | Mouse | Novus Biologicals | NBP2-33200AF750 |  |
| Fig. 5–8 IF |  | AF488 | CD7 | membrane | EPR4242 | 5 | Rabbit | Abcam | ab199022 |  |
|  |  | AF555 | CD3 | membrane | EP4426 | 1 | Rabbit | Abcam | ab208514 |  |
|  |  | AF647 | CD5 | membrane | EP2952 | 5 | Rabbit | Abcam | ab285357 |  |
| Fig. 6 BF |  | none | CD3 | membrane | LN10 | Ready to use | mouse | Leica Biosystems | PA0553 |  |
|  |  | none | CD5 | membrane | 4C7 | Ready to use | mouse | Leica Biosystems | PA0168 |  |
|  |  | none | CD7 | membrane | LP15 | Ready to use | mouse | Leica Biosystems | PA0266 |  |
| Supplemental Fig. 2a |  | none | CD4 | membrane | 4B12 | Ready to use | mouse | Leica Biosystems | PA0427 |  |
|  |  | none | PD-1 | membrane | NAT-105 | 5 | mouse | Abcam | ab52587 |  |
|  |  | none | Ki-67 | nucleus | MM1 | Ready to use | mouse | Leica Biosystems | PA0118 |  |
|  |  | none | PD-L1 | membrane | 73-10 | Ready to use | Rabbit | Leica Biosystems | PA0832 |  |
|  |  | none | CD3 | membrane | LN10 | Ready to use | mouse | Leica Biosystems | PA0553 |  |
|  |  | none | Cytokeratin | cytoplasm | AE1-AE3 | Ready to use | mouse | Leica Biosystems | PA0094 |  |
|  |  | none | CD8 | membrane | 4B11 | Ready to use | mouse | Leica Biosystems | PA0183 |  |
|  |  | none | CD68 | cytoplasm | 514H12 | Ready to use | mouse | Leica Biosystems | PA0273 |  |
| Supplemental Fig. 2b |  | none | CD45RO | membrane | UCHL1 | Ready to use | mouse | Leica Biosystems | PA0146 |  |
|  |  | none | CD3 | membrane | LN10 | Ready to use | mouse | Leica Biosystems | PA0553 |  |
|  |  | none | PD-1 | membrane | NAT-105 | 5 | mouse | Abcam | ab52587 |  |
|  |  | none | SOX10 | nucleus | SP267 | 1 | mouse | Abcam | ab227680 |  |
|  |  | none | Cytokeratin | cytoplasm | AE1-AE3 | Ready to use | mouse | Leica Biosystems | PA0094 |  |
| Supplemental Fig. 2c |  | none | CD4 | membrane | 4B12 | Ready to use | mouse | Leica Biosystems | PA0427 |  |
|  |  | none | Foxp3 | nucleus | D2W8E | 3 | Rabbit | Cell Signaling Technology | 98377S |  |
|  |  | none | CD20 | membrane | L26 | Ready to use | mouse | Leica Biosystems | PA0200 |  |
|  |  | none | CD8 | membrane | 4B11 | Ready to use | mouse | Leica Biosystems | PA0183 |  |
|  |  | none | CD68 | cytoplasm | 514H12 | Ready to use | mouse | Leica Biosystems | PA0273 |  |
|  |  | none | Cytokeratin | cytoplasm | AE1-AE3 | Ready to use | mouse | Leica Biosystems | PA0094 |  |

**Supplemental Figure 1. Hyper Spectral Imaging System.**

(a) Schematic outline of the hyperspectral imaging system. A custom-made multispectral scanner facilitates the acquisition of data across 135 channels. Four line-shaped beam profiles are employed for excitation, with each laser sequentially scanning the stained sample. The resultant fluorescence is analyzed using a spectrometer, with data captured using an image sensor. (b) Representation of spatio-spectral signals on the image sensor, where spectroscopic fluorescence is captured, displaying spatial information on the horizontal axis and spectroscopic information on the vertical axis. (c) Description of the spatial sweep spectroscopic hyperspectral imaging prototype system.

**Supplemental Figure 2. Spectral Unmixing Algorithms.**

(a) Schematic of spectral optimization using NMF. The measured spectral cube data A (Row: pixel, Column: fluorescent spectra) is dimensionally reduced to the fluorescence spectra element H (Row: wavelength, Column: fluorescent intensity) and the standardized intensity element W (Row: fluorophore species, Column: pixel) by optimizing two elements to minimize residuals Δ without allowing negative values. (b) Conceptual diagram for acquiring reference spectra for standardized intensity. A dispersed sample at a known concentration is measured on a sensor with a defined volume, expressing the number of fluorescent antibodies present in a voxel corresponding to a sensor pixel as [the number of molecules in sensor 1 pix = M × ST × Avogadro number], utilizing the concentration (M) of each dye and the area of one-pixel sensor (S) and the thickness of slide glass (T).

**Supplemental Figure 3. Antibody Validation using Immunohistochemistry (IHC).**

IHC/DAB staining for each antibody utilized in Figure 4, demonstrating their specificity and staining patterns.

**Supplemental Figure 4. Differences in Marker Expression Distribution Between Samples.**

Density scatter plot analysis of CD3, CD5, and CD7 expression at the single-cell level, derived from image cytometry analysis, showcasing the variance between samples. (a) CD3 and CD5 expression between Sample 1 (Control) and Sample 26, with the average number of CD3 and CD5 antibodies per voxel for each cell plotted. The analysis is visualized with the x-axis representing CD3 and the y-axis representing CD5 antibody counts per voxel. Red lines mark thresholds set at 7 × 100 for CD3 and 2 × 100 for CD5, indicating specific expression levels considered significant.

(b) CD3 and CD7 expression in the same samples, with axes representing averaged antibody counts per voxel for CD3 and CD7, respectively. The delineation thresholds are illustrated by red lines, set at 7 × 100 for CD3 and 2 × 10^1 for CD7, highlighting the criteria used to differentiate expression intensities within the sampled cells.

**Supplemental Figure 5. Non-Negative Matrix Factorization (NMF)**

Overview of the jNMF algorithm, illustrating the decomposition of matrices X1, X2, ..., Xm (purple) into a common factor matrix W (red) and m factor matrices H1, H2, ..., Hm (blue) as feature vectors.
