## Supplemental figures for "Spatial Clustering Analysis with Spectral Imaging-based Single-Step Multiplex Immunofluorescence (SISS-mIF) for Assisting Histological Diagnosis"

### Slide 1
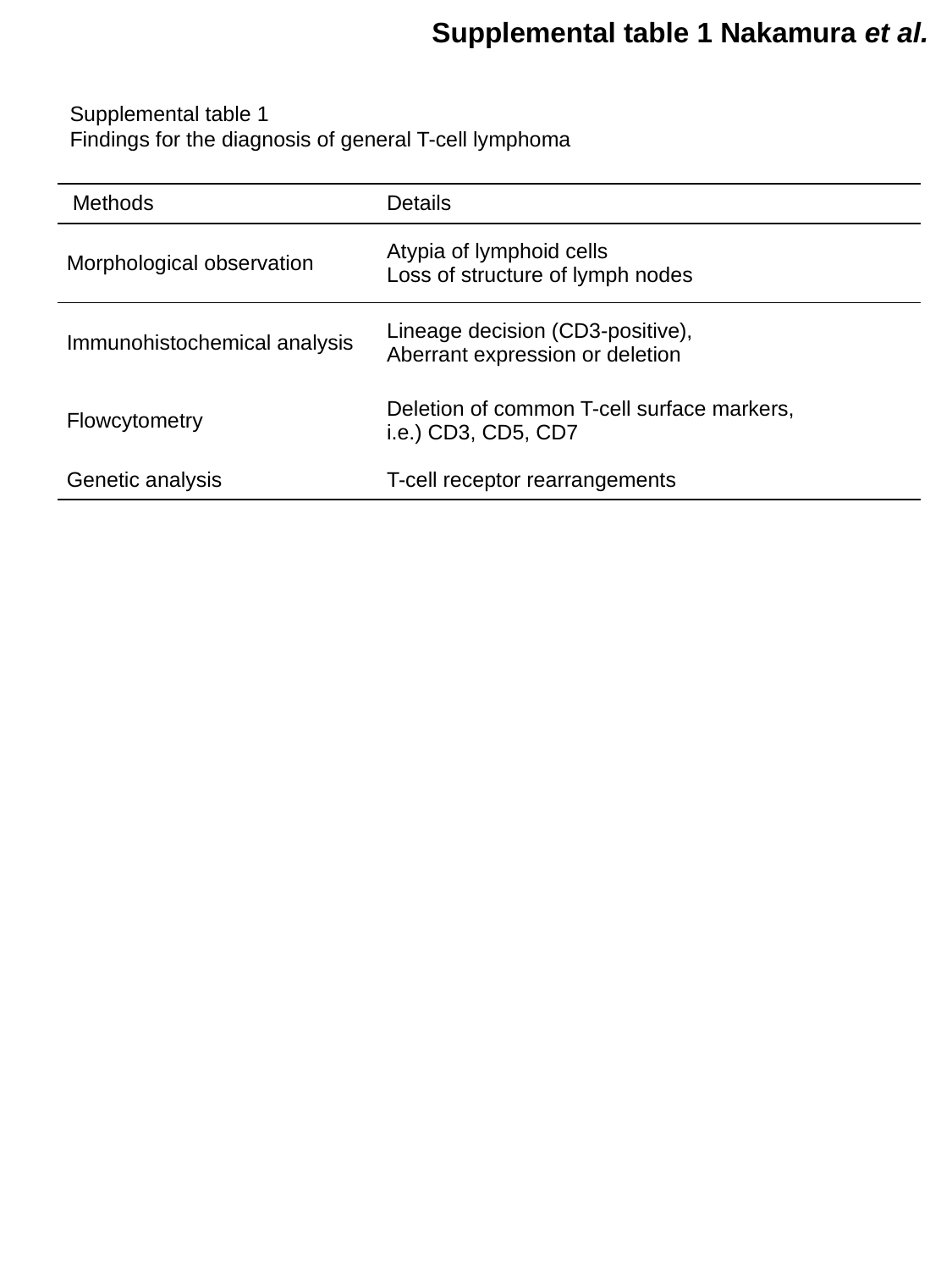

Supplemental table 1 Nakamura et al.
Supplemental table 1
Findings for the diagnosis of general T-cell lymphoma
| Methods | Details |
| --- | --- |
| Morphological observation | Atypia of lymphoid cells Loss of structure of lymph nodes |
| Immunohistochemical analysis | Lineage decision (CD3-positive), Aberrant expression or deletion |
| Flowcytometry | Deletion of common T-cell surface markers, i.e.) CD3, CD5, CD7 |
| Genetic analysis | T-cell receptor rearrangements |

### Slide 2
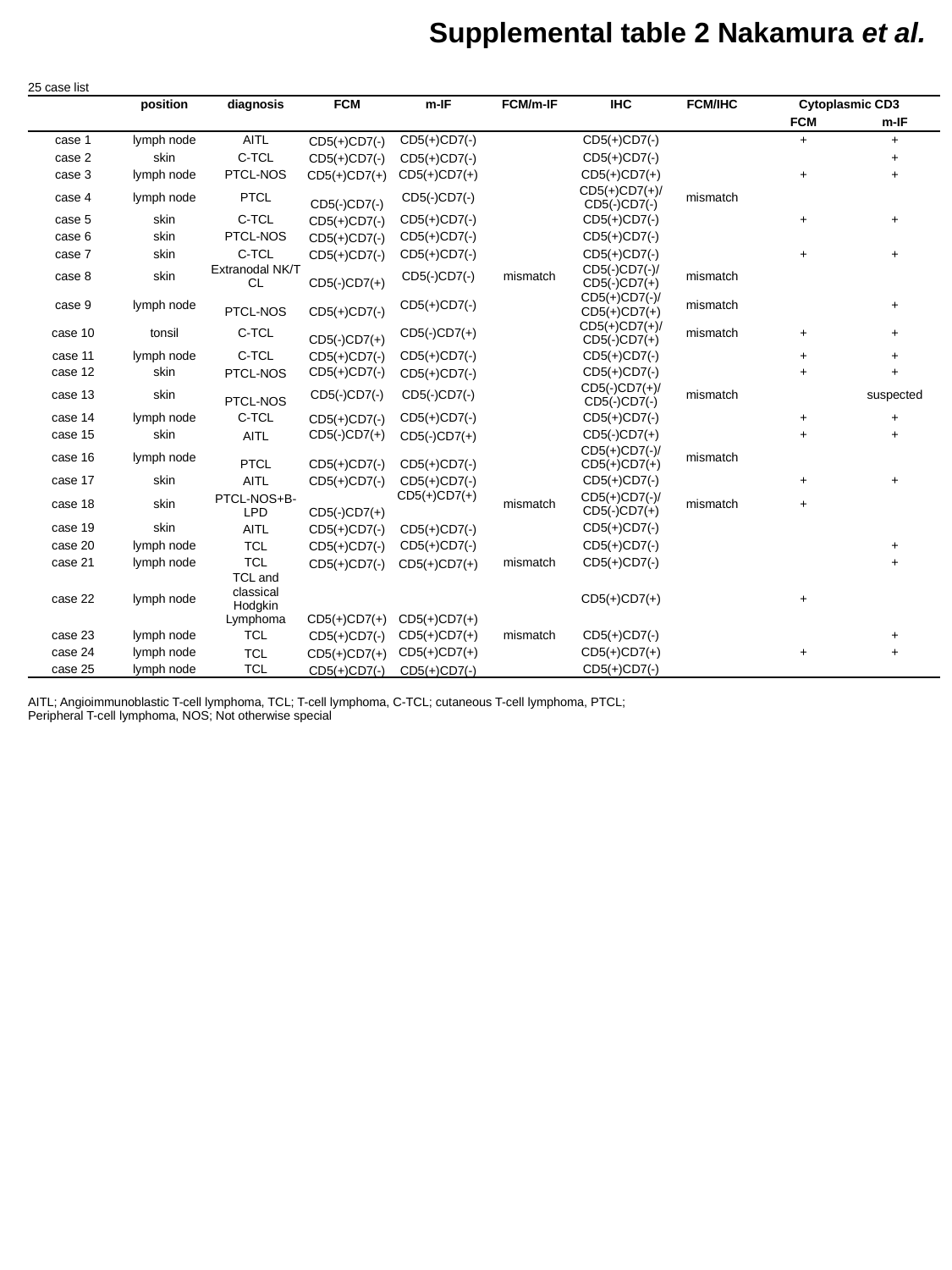

Supplemental table 2 Nakamura et al.
| 25 case list | | | | | | | | | |
| --- | --- | --- | --- | --- | --- | --- | --- | --- | --- |
| | position | diagnosis | FCM | m-IF | FCM/m-IF | IHC | FCM/IHC | Cytoplasmic CD3 | |
| | | | | | | | | FCM | m-IF |
| case 1 | lymph node | AITL | CD5(+)CD7(-) | CD5(+)CD7(-) | | CD5(+)CD7(-) | | + | + |
| case 2 | skin | C-TCL | CD5(+)CD7(-) | CD5(+)CD7(-) | | CD5(+)CD7(-) | | | + |
| case 3 | lymph node | PTCL-NOS | CD5(+)CD7(+) | CD5(+)CD7(+) | | CD5(+)CD7(+) | | + | + |
| case 4 | lymph node | PTCL | CD5(-)CD7(-) | CD5(-)CD7(-) | | CD5(+)CD7(+)/CD5(-)CD7(-) | mismatch | | |
| case 5 | skin | C-TCL | CD5(+)CD7(-) | CD5(+)CD7(-) | | CD5(+)CD7(-) | | + | + |
| case 6 | skin | PTCL-NOS | CD5(+)CD7(-) | CD5(+)CD7(-) | | CD5(+)CD7(-) | | | |
| case 7 | skin | C-TCL | CD5(+)CD7(-) | CD5(+)CD7(-) | | CD5(+)CD7(-) | | + | + |
| case 8 | skin | Extranodal NK/T CL | CD5(-)CD7(+) | CD5(-)CD7(-) | mismatch | CD5(-)CD7(-)/CD5(-)CD7(+) | mismatch | | |
| case 9 | lymph node | PTCL-NOS | CD5(+)CD7(-) | CD5(+)CD7(-) | | CD5(+)CD7(-)/CD5(+)CD7(+) | mismatch | | + |
| case 10 | tonsil | C-TCL | CD5(-)CD7(+) | CD5(-)CD7(+) | | CD5(+)CD7(+)/CD5(-)CD7(+) | mismatch | + | + |
| case 11 | lymph node | C-TCL | CD5(+)CD7(-) | CD5(+)CD7(-) | | CD5(+)CD7(-) | | + | + |
| case 12 | skin | PTCL-NOS | CD5(+)CD7(-) | CD5(+)CD7(-) | | CD5(+)CD7(-) | | + | + |
| case 13 | skin | PTCL-NOS | CD5(-)CD7(-) | CD5(-)CD7(-) | | CD5(-)CD7(+)/CD5(-)CD7(-) | mismatch | | suspected |
| case 14 | lymph node | C-TCL | CD5(+)CD7(-) | CD5(+)CD7(-) | | CD5(+)CD7(-) | | + | + |
| case 15 | skin | AITL | CD5(-)CD7(+) | CD5(-)CD7(+) | | CD5(-)CD7(+) | | + | + |
| case 16 | lymph node | PTCL | CD5(+)CD7(-) | CD5(+)CD7(-) | | CD5(+)CD7(-)/CD5(+)CD7(+) | mismatch | | |
| case 17 | skin | AITL | CD5(+)CD7(-) | CD5(+)CD7(-) | | CD5(+)CD7(-) | | + | + |
| case 18 | skin | PTCL-NOS+B-LPD | CD5(-)CD7(+) | CD5(+)CD7(+) | mismatch | CD5(+)CD7(-)/CD5(-)CD7(+) | mismatch | + | |
| case 19 | skin | AITL | CD5(+)CD7(-) | CD5(+)CD7(-) | | CD5(+)CD7(-) | | | |
| case 20 | lymph node | TCL | CD5(+)CD7(-) | CD5(+)CD7(-) | | CD5(+)CD7(-) | | | + |
| case 21 | lymph node | TCL | CD5(+)CD7(-) | CD5(+)CD7(+) | mismatch | CD5(+)CD7(-) | | | + |
| case 22 | lymph node | TCL and classical Hodgkin Lymphoma | CD5(+)CD7(+) | CD5(+)CD7(+) | | CD5(+)CD7(+) | | + | |
| case 23 | lymph node | TCL | CD5(+)CD7(-) | CD5(+)CD7(+) | mismatch | CD5(+)CD7(-) | | | + |
| case 24 | lymph node | TCL | CD5(+)CD7(+) | CD5(+)CD7(+) | | CD5(+)CD7(+) | | + | + |
| case 25 | lymph node | TCL | CD5(+)CD7(-) | CD5(+)CD7(-) | | CD5(+)CD7(-) | | | |
| AITL; Angioimmunoblastic T-cell lymphoma, TCL; T-cell lymphoma, C-TCL; cutaneous T-cell lymphoma, PTCL; Peripheral T-cell lymphoma, NOS; Not otherwise special | | | | | | | | | |

### Slide 3
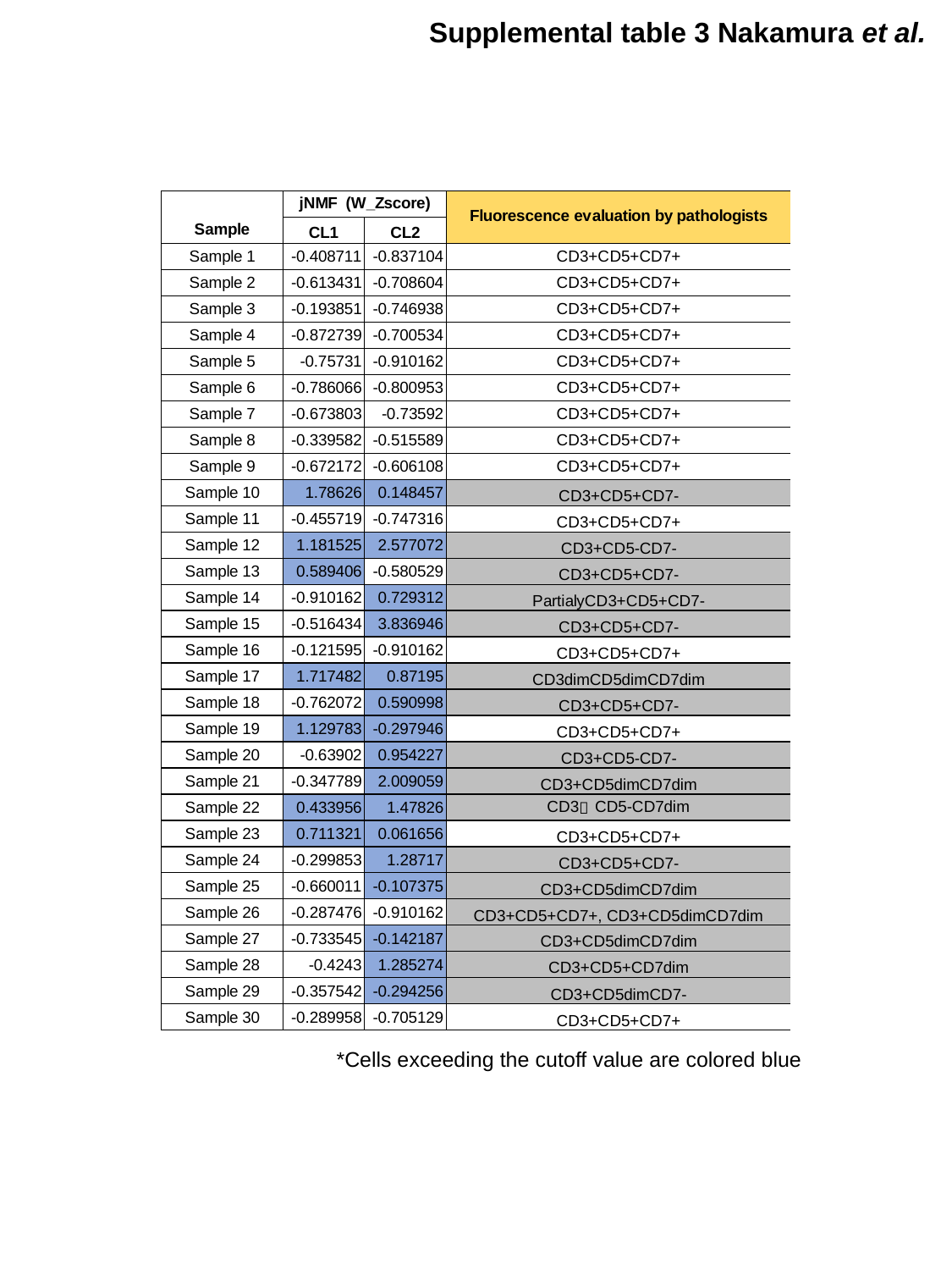

Supplemental table 3 Nakamura et al.
*Cells exceeding the cutoff value are colored blue

### Slide 4
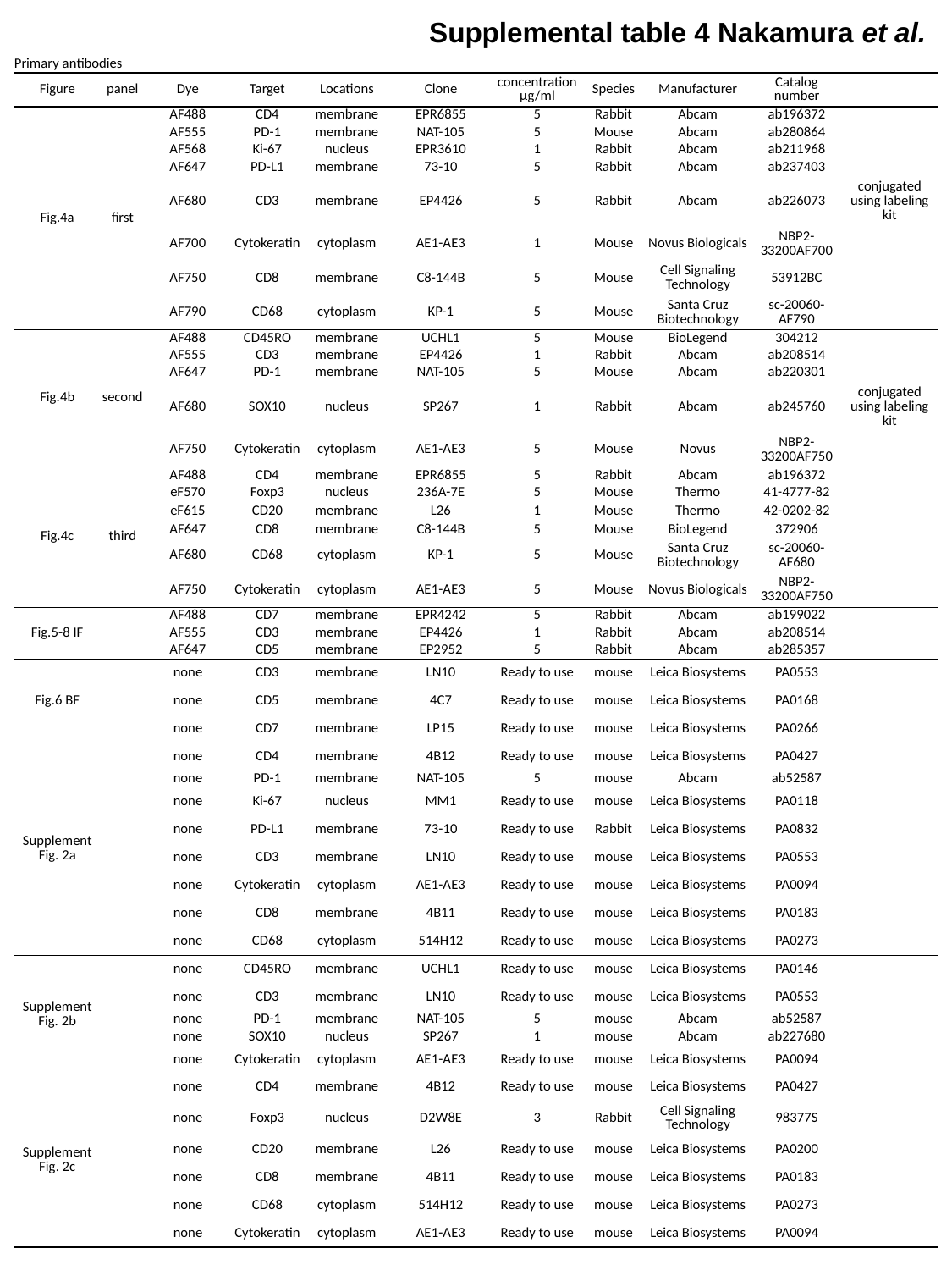

Supplemental table 4 Nakamura et al.
| Primary antibodies | | | | | | | | | | |
| --- | --- | --- | --- | --- | --- | --- | --- | --- | --- | --- |
| Figure | panel | Dye | Target | Locations | Clone | concentration μg/ml | Species | Manufacturer | Catalog number | |
| Fig.4a | first | AF488 | CD4 | membrane | EPR6855 | 5 | Rabbit | Abcam | ab196372 | |
| | | AF555 | PD-1 | membrane | NAT-105 | 5 | Mouse | Abcam | ab280864 | |
| | | AF568 | Ki-67 | nucleus | EPR3610 | 1 | Rabbit | Abcam | ab211968 | |
| | | AF647 | PD-L1 | membrane | 73-10 | 5 | Rabbit | Abcam | ab237403 | |
| | | AF680 | CD3 | membrane | EP4426 | 5 | Rabbit | Abcam | ab226073 | conjugated using labeling kit |
| | | AF700 | Cytokeratin | cytoplasm | AE1-AE3 | 1 | Mouse | Novus Biologicals | NBP2-33200AF700 | |
| | | AF750 | CD8 | membrane | C8-144B | 5 | Mouse | Cell Signaling Technology | 53912BC | |
| | | AF790 | CD68 | cytoplasm | KP-1 | 5 | Mouse | Santa Cruz Biotechnology | sc-20060-AF790 | |
| Fig.4b | second | AF488 | CD45RO | membrane | UCHL1 | 5 | Mouse | BioLegend | 304212 | |
| | | AF555 | CD3 | membrane | EP4426 | 1 | Rabbit | Abcam | ab208514 | |
| | | AF647 | PD-1 | membrane | NAT-105 | 5 | Mouse | Abcam | ab220301 | |
| | | AF680 | SOX10 | nucleus | SP267 | 1 | Rabbit | Abcam | ab245760 | conjugated using labeling kit |
| | | AF750 | Cytokeratin | cytoplasm | AE1-AE3 | 5 | Mouse | Novus | NBP2-33200AF750 | |
| Fig.4c | third | AF488 | CD4 | membrane | EPR6855 | 5 | Rabbit | Abcam | ab196372 | |
| | | eF570 | Foxp3 | nucleus | 236A-7E | 5 | Mouse | Thermo | 41-4777-82 | |
| | | eF615 | CD20 | membrane | L26 | 1 | Mouse | Thermo | 42-0202-82 | |
| | | AF647 | CD8 | membrane | C8-144B | 5 | Mouse | BioLegend | 372906 | |
| | | AF680 | CD68 | cytoplasm | KP-1 | 5 | Mouse | Santa Cruz Biotechnology | sc-20060-AF680 | |
| | | AF750 | Cytokeratin | cytoplasm | AE1-AE3 | 5 | Mouse | Novus Biologicals | NBP2-33200AF750 | |
| Fig.5-8 IF | | AF488 | CD7 | membrane | EPR4242 | 5 | Rabbit | Abcam | ab199022 | |
| | | AF555 | CD3 | membrane | EP4426 | 1 | Rabbit | Abcam | ab208514 | |
| | | AF647 | CD5 | membrane | EP2952 | 5 | Rabbit | Abcam | ab285357 | |
| Fig.6 BF | | none | CD3 | membrane | LN10 | Ready to use | mouse | Leica Biosystems | PA0553 | |
| | | none | CD5 | membrane | 4C7 | Ready to use | mouse | Leica Biosystems | PA0168 | |
| | | none | CD7 | membrane | LP15 | Ready to use | mouse | Leica Biosystems | PA0266 | |
| Supplement Fig. 2a | | none | CD4 | membrane | 4B12 | Ready to use | mouse | Leica Biosystems | PA0427 | |
| | | none | PD-1 | membrane | NAT-105 | 5 | mouse | Abcam | ab52587 | |
| | | none | Ki-67 | nucleus | MM1 | Ready to use | mouse | Leica Biosystems | PA0118 | |
| | | none | PD-L1 | membrane | 73-10 | Ready to use | Rabbit | Leica Biosystems | PA0832 | |
| | | none | CD3 | membrane | LN10 | Ready to use | mouse | Leica Biosystems | PA0553 | |
| | | none | Cytokeratin | cytoplasm | AE1-AE3 | Ready to use | mouse | Leica Biosystems | PA0094 | |
| | | none | CD8 | membrane | 4B11 | Ready to use | mouse | Leica Biosystems | PA0183 | |
| | | none | CD68 | cytoplasm | 514H12 | Ready to use | mouse | Leica Biosystems | PA0273 | |
| Supplement Fig. 2b | | none | CD45RO | membrane | UCHL1 | Ready to use | mouse | Leica Biosystems | PA0146 | |
| | | none | CD3 | membrane | LN10 | Ready to use | mouse | Leica Biosystems | PA0553 | |
| | | none | PD-1 | membrane | NAT-105 | 5 | mouse | Abcam | ab52587 | |
| | | none | SOX10 | nucleus | SP267 | 1 | mouse | Abcam | ab227680 | |
| | | none | Cytokeratin | cytoplasm | AE1-AE3 | Ready to use | mouse | Leica Biosystems | PA0094 | |
| Supplement Fig. 2c | | none | CD4 | membrane | 4B12 | Ready to use | mouse | Leica Biosystems | PA0427 | |
| | | none | Foxp3 | nucleus | D2W8E | 3 | Rabbit | Cell Signaling Technology | 98377S | |
| | | none | CD20 | membrane | L26 | Ready to use | mouse | Leica Biosystems | PA0200 | |
| | | none | CD8 | membrane | 4B11 | Ready to use | mouse | Leica Biosystems | PA0183 | |
| | | none | CD68 | cytoplasm | 514H12 | Ready to use | mouse | Leica Biosystems | PA0273 | |
| | | none | Cytokeratin | cytoplasm | AE1-AE3 | Ready to use | mouse | Leica Biosystems | PA0094 | |

### Slide 5
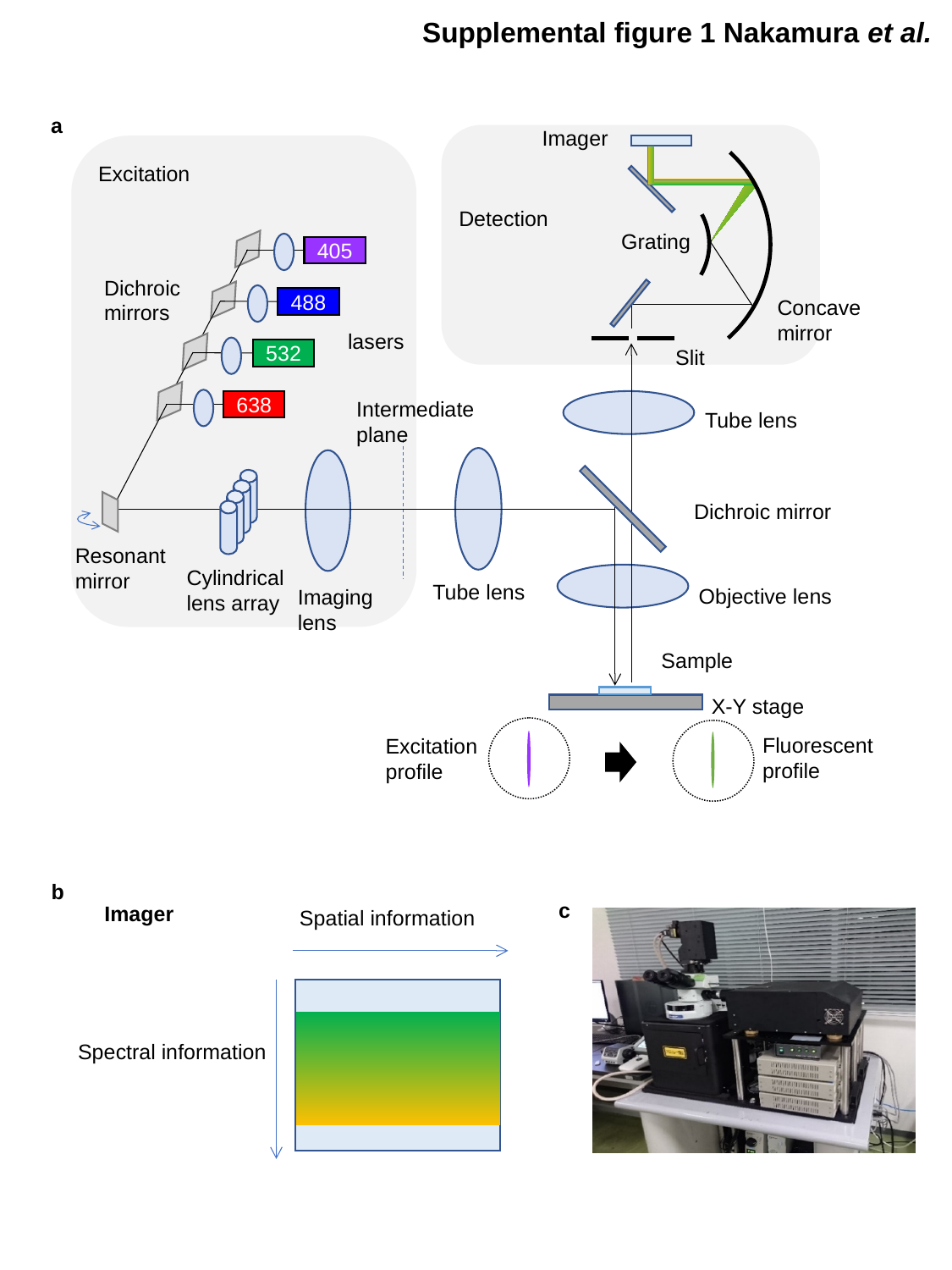

Supplemental figure 1 Nakamura et al.
a
Imager
Excitation
Detection
Grating
405
Dichroic mirrors
488
Concave
mirror
lasers
Slit
532
Intermediate plane
638
Tube lens
Dichroic mirror
Resonant mirror
Cylindrical lens array
Tube lens
Objective lens
Imaging lens
Sample
X-Y stage
Fluorescent profile
Excitation profile
b
c
Imager
Spatial information
Spectral information

### Slide 6
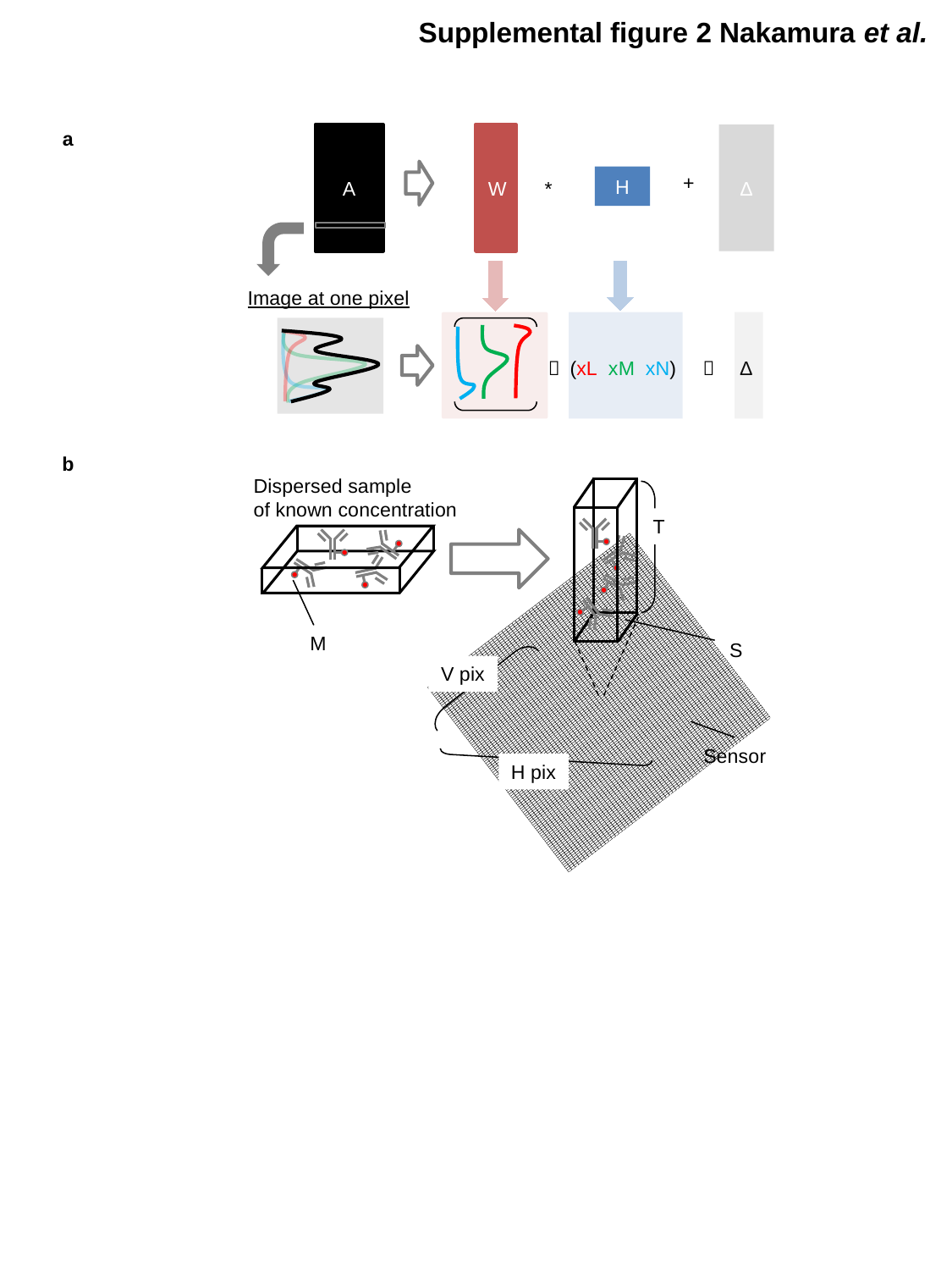

Supplemental figure 2 Nakamura et al.
a
A
W
Δ
+
H
*
Image at one pixel
＊ (xL xM xN)
＋ Δ
b
Dispersed sample
of known concentration
T
M
S
V pix
Sensor
H pix

### Slide 7
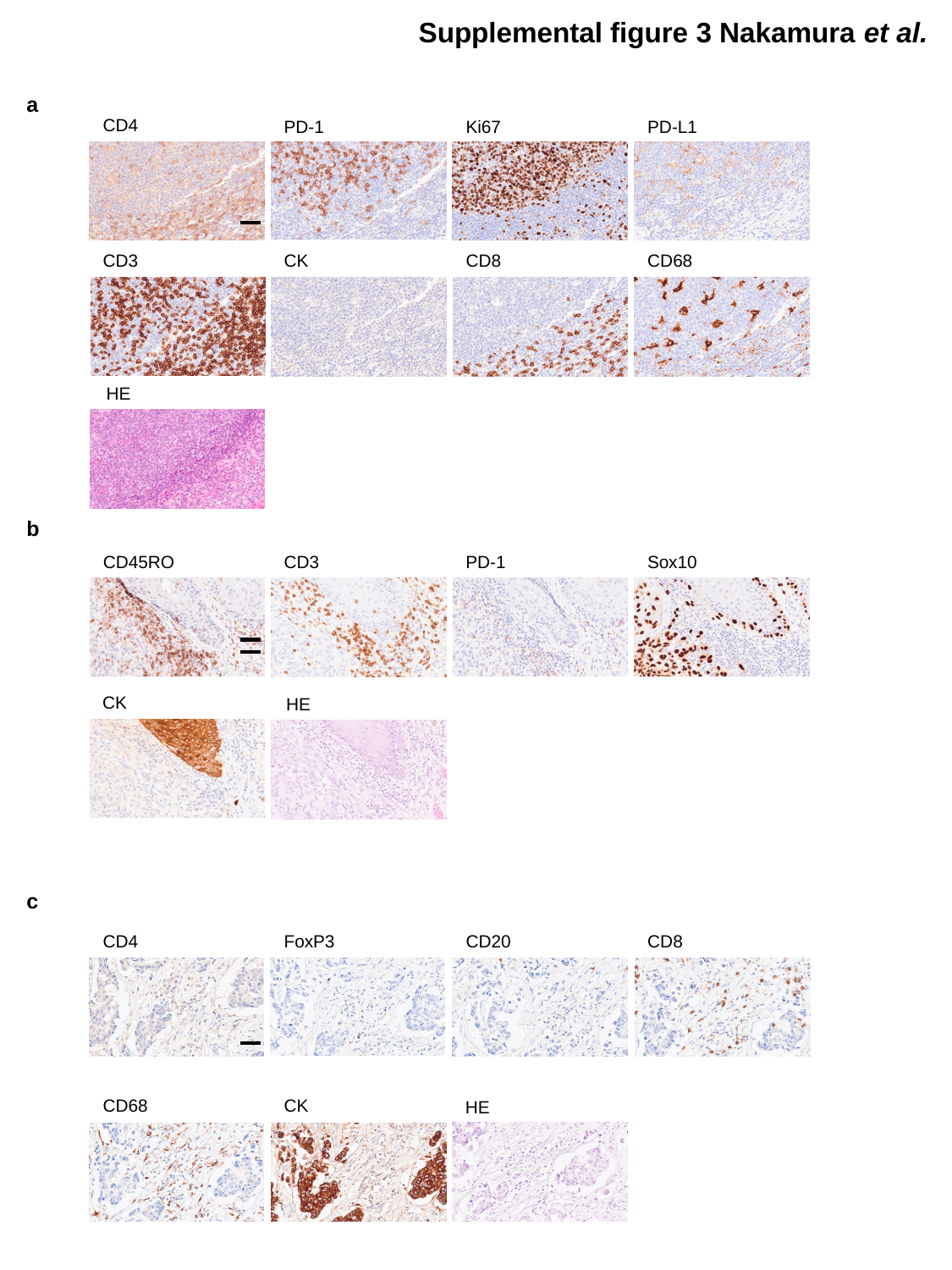

Supplemental figure 3 Nakamura et al.
a
CD4
PD-1
Ki67
PD-L1
CD3
CK
CD8
CD68
HE
b
CD45RO
CD3
PD-1
Sox10
CK
HE
c
CD4
FoxP3
CD20
CD8
CD68
CK
HE

### Slide 8
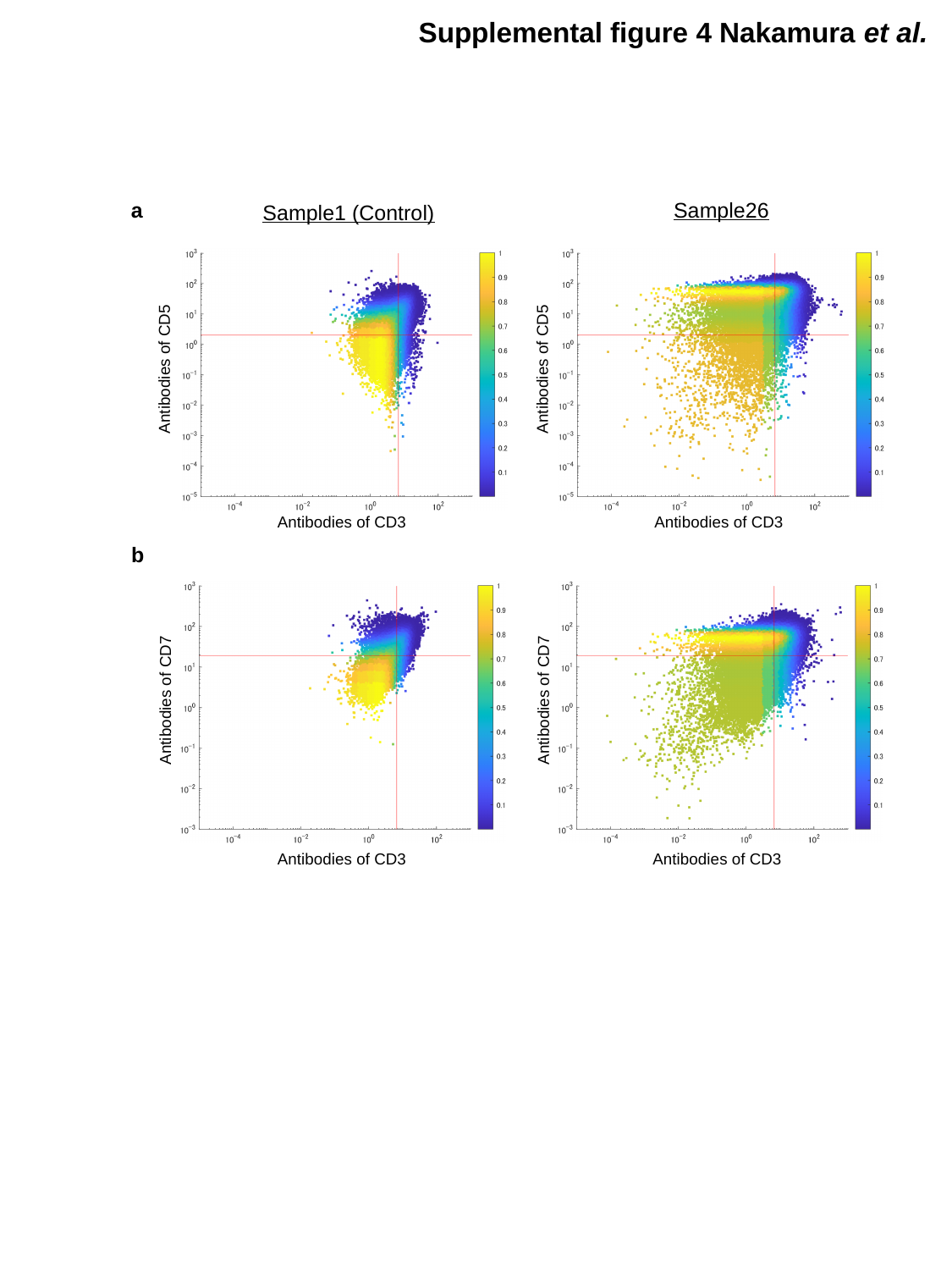

Supplemental figure 4 Nakamura et al.
a
Sample26
Sample1 (Control)
Antibodies of CD5
Antibodies of CD5
Antibodies of CD3
Antibodies of CD3
b
Antibodies of CD7
Antibodies of CD7
Antibodies of CD3
Antibodies of CD3

### Slide 9
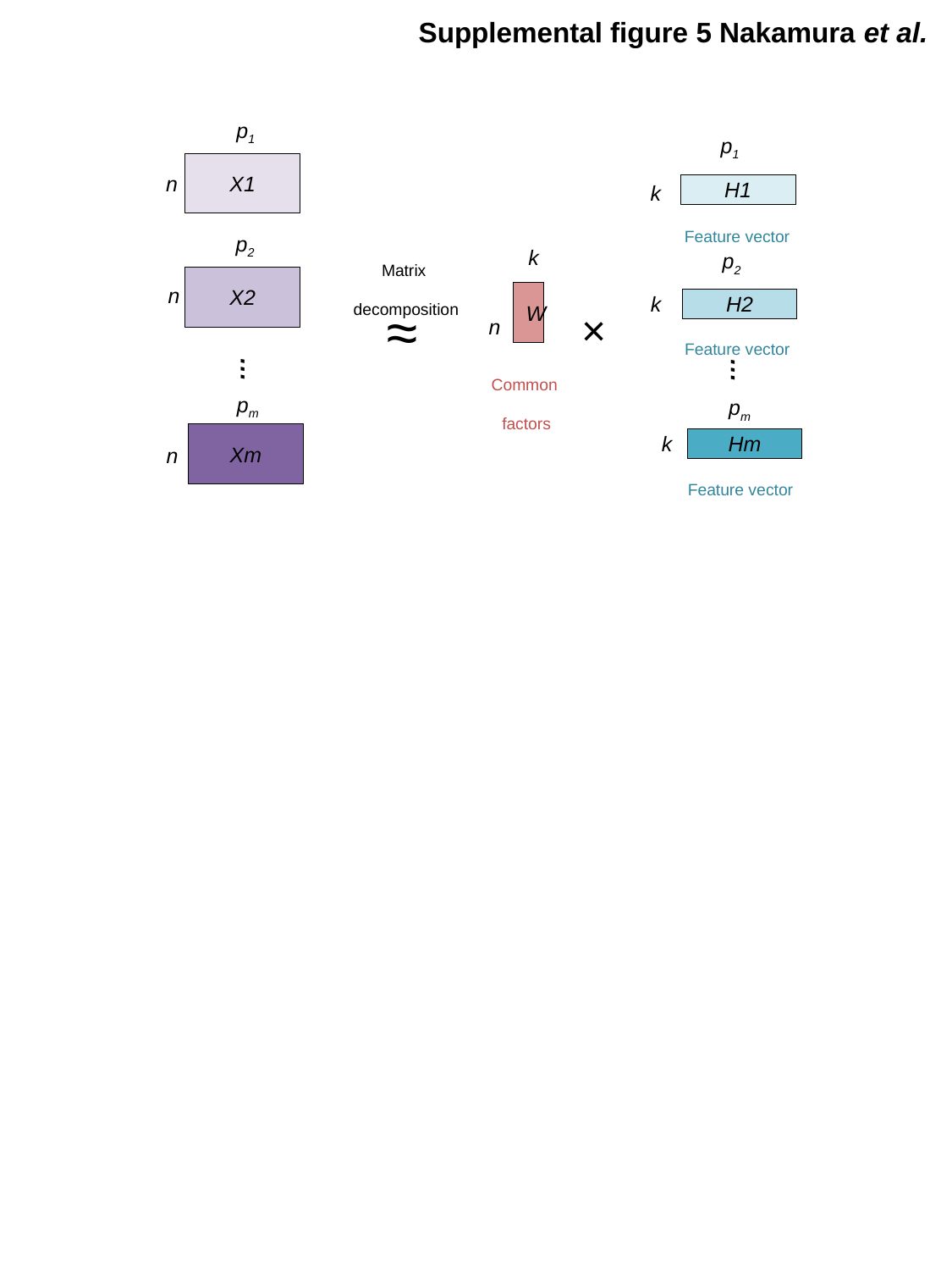

Supplemental figure 5 Nakamura et al.
p_1_
p_1_
H1
X1
n
k
_Feature vector_
p_2_
_Matrix_
 _decomposition_
k
p_2_
H2
X2
n
W
k
≈
×
n
_Feature vector_
…
_Common_
_factors_
…
p_m_
Hm
p_m_
Xm
k
n
_Feature vector_
